# Axiomatic-deductive theory of competition of complete competitors: coexistence, exclusion and neutrality in one model

**DOI:** 10.1101/2020.07.22.215533

**Authors:** Lev V. Kalmykov, Vyacheslav L. Kalmykov

## Abstract

**Background:** The long-standing contradiction between formulations of the competitive exclusion principle and natural diversity of trophically similar species is known as the biodiversity paradox. Earlier we found that coexistence of complete competitors is possible despite 100% difference in competitiveness, but only under certain conditions – at their moderate propagation and at the particular initial location of individuals. Here we verify a hypothesis that completely competing species with aggressive propagation may coexist with less than 100% difference in competitiveness regardless of random initial location of competing individuals in ecosystem.

**Methods:** We investigate a role of competitiveness differences in coexistence of two completely competing species by individual-based modeling based on a transparent artificial intelligence. We propose and investigate an individual-based model of ecosystem dynamics supplemented by a probabilistic determination of the competitiveness of competing individuals without cooperative effects and with cooperative effects based on the numerical superiority of individuals of the species.

**Results:** We have found that two aggressively propagating complete competitors can stably coexist, despite one species has some advantage in competitiveness over the other and all other characteristics of the species are equal. The found competitive coexistence occurred regardless of the initial random location of individuals in the ecosystem. When colonization of a free habitat started from a single individual of each species, then the complete competitors coexisted up to 31% of their difference in competitiveness. And when on initial stage half of the territory was probabilistically occupied, the complete competitors coexisted up to 22% of their difference in competitiveness. In the experiments with cooperative dependence on the numerical superiority of individuals of the species complete competitors stably co-existed despite 10% difference in basic competitiveness.

**Discussion:** The results additionally support our earlier reformulation of the competitive exclusion principle. Besides that, we revealed classical cases of competitive exclusion and “neutrality”. Our approach unifies models of competitive exclusion (“niche”), neutrality and coexistence of complete competitors in one theory. Our individual-based modeling of a complex system based on a transparent artificial intelligence opens up great prospects for a variety of theoretical and applied fields.

## 1 INTRODUCTION

### 1.1 The central issue in theoretical ecology

Mechanisms of interspecific competition underlying the theory of evolution still remain insufficiently understood (den Boer 1986, Tilman 1987, Sommer 1999, Whitfield 2002, Clark 2009, Clark 2012). The conceptual basis of the Darwinian natural selection is the statement about the survival of the fittest. A key tool for implementing natural selection is the principle of competitive exclusion - complete competitors cannot coexist if they occupy precisely the same resource niche (Gause 1934, Gause and Witt 1935, Hardin 1959, Hardin 1960). In theory, according to the competitive exclusion principle, complete competitors cannot coexist, but in practice, there are many examples of the trophically similar species coexistence: tropical rainforest, coral reefs, plankton communities. This contradiction is known as the biodiversity paradox (Hutchinson 1961, Sommer 1999). Resolving the diversity paradox became the central issue in theoretical ecology (Lehman and Tilman 1997). On the terms – the “complete competitors” means that competitors are identical in consumption, i.e. they consume identical resources in the same way. At the same time, their competitiveness as and the methods of competition may differ. We define competitiveness as a probability of a species to propagate by occupation of a free microhabitat by its individual in direct local interspecific competition. The urgent tasks of biodiversity conservation became the additional motivation of the long-standing biodiversity debates. As diversity of species of similar trophic level is the fact, then the problem of solving the paradox is reduced to verification of the formulations of the competitive exclusion principle. Understanding the mechanisms of interspecific competition is complicated, among other things, by the crisis in the field of classical rationality, which has been going on since the beginning of the last century. This crisis arose due to the emergence of fundamentally new discoveries that did not fit into the existing system of knowledge and problems associated with the need to understand especially complex systems. The ideals of a clear transparent understanding of the causal mechanisms of the phenomena under study began to be replaced by opaque approaches that exaggerate the role of uncertainty, ambiguity and relativity of knowledge, allowing the absence of cause-and-effect relationships in the phenomena under study. Black box models, which can predict but cannot explain anything, have become dominant.

We believe that the general confusion over understanding the mechanisms of interspecific competition stems from the four sources:

1. The first source is the impossibility of thorough experimental testing of hypothetical mechanisms that implement the principle of competitive exclusion, which was explained by (Hardin 1960). An experimental verification of the validity of the principle of competitive exclusion is impossible, because we will always suspect that some experimental factor was not taken into account – “There are many who have supposed that the principle is one that can be proved or disproved by empirical facts, among them Gauze himself. Nothing could be farther from the truth. The “truth” of the principle is and can be established only by theory, not being subject to proof or disproof by facts, as ordinarily understood” (Hardin 1960).
2. The second source is the opacity of mathematical models, which complicates interpretation of their results (Tilman 1987, Kalmykov and Kalmykov 2016) drew attention to the fact that ecologists investigate interspecific competition phenomenologically, rather than mechanistically – “most ecologists have studied competition by asking if an increase in the density of one species leads to a decrease in the density of another, without asking how this might occur” (Tilman 1987). The nontransparent phenomenological models show what happens with the modelled object on a macro-level but does not show how it happens on a micro-level of individuals (Kalmykov and Kalmykov 2016). They describe some empirical observations, but have no foundations in underlying mechanisms or first principles. It makes difficult a prediction, generation of new knowledge and creation of new technologies. The most well-known model of interspecific competition is the Lotka-Volterra model. The opacity of the Lotka-Volterra model forces researchers to come up with interpretations of the underlying mechanisms without getting a complete clarity regarding the mechanisms being studied. This model is phenomenological (Tilman 1987, Huston et al. 1988a, Kalmykov and Kalmykov 2013, Kalmykov and Kalmykov 2016) because, despite the fact that it is deterministic, it does not model local interactions. This model predicts stable coexistence of two similar species when, for both species, an interspecific competition is weaker than intraspecific one. This interpretation follows directly from the model. However, the further reinterpretation in the form of the competitive exclusion principle has no rigorous justification under itself. The unified neutral theory of biodiversity and biogeography (neutral theory or UNTB) was proposed as an attempt to solve the biodiversity paradox (Hubbell 2001). But the neutral theory proposed to replace the study of mechanisms of interspecific interactions by statistical predictions of species presence-absence: “We no longer need better theories of species coexistence; we need better theories for species presence-absence, relative abundance and persistence times in communities that can be confronted with real data. In short, it is long past time for us to get over our myopic preoccupation with coexistence” (Hubbell 2001). This way is based on the assumptions which are clearly not true as “no ecologist believes that species are identical in reality” (Doncaster 2009). Unreality of the hypothesis of ecological equivalence is obvious and for Hubbell with colleagues, as they assert that *“the real world is not neutral”* (Rosindell et al. 2012). The long-standing debates on the biodiversity paradox “competitive exclusion principle versus natural biodiversity” has been substituted for theoretical debates *“neutrality versus the niche”* (Whitfield 2002, Kalmykov and Kalmykov 2016). There is even an attempt “due to an objective limitation of nature” to introduce in ecology principle of quantum environmental uncertainty, which based on non-transparent models of quantum mechanics. According to this principle it is fundamentally impossible to prove the validity of one of two hypotheses which are clearly contradictory - they “are simultaneously true and false in the same measure, because the only feasible option to keep the functional stability of ecosystems is a wave-like combination of both options…” (Rodríguez et al. 2015, Rodríguez et al. 2016). Accusations of nature regarding its “*objective limitation*” (Rodríguez et al. 2016) we consider as a consequence of limitations of opaque mathematical models used by researchers, but not of nature itself. To overcome these difficulties, it is necessary to use transparent mathematical models that allow modeling global behavior from local rules (Huston et al. 1988b, Silvertown et al. 1992, Kalmykov and Kalmykov 2016).
3. The third source of difficulties is the multifactorial nature of the interspecific competition. Combinations of multiple diversity of ecological resources and environmental conditions, of intrinsic resource preferences and functional traits of competing species generates a lot of mechanisms which promote coexistence (Hutchinson 1961, Hastings 1980, Palmer 1994, Chesson 2000, Wellborn 2002, Dollhopf et al. 2003, Nowak 2006, Bennett and Bever 2009): trade-offs; cooperative interactions between competing species; genetic heterogeneity of populations; sexual reproduction, which increases the genetic heterogeneity of populations; excess ecosystem resources; heterogeneity of habitat; immigration, emigration, predation, parasitism, herbivory, diseases and other problems of the population; variability in environmental conditions; ontogenetic differences in competitiveness, in fertility rates, in the features of habitat regeneration and in the environmental requirements of competing species. These mechanisms complicate testing the hypotheses about conditions of implementing the competitive exclusion principle. An example of difficulties arising from the existence of a large number of coexistence mechanisms is the commentary of Pastor et al. (Pásztor et al. 2020) to the article McPeek (McPeek 2019). Commentators are argued that the mechanism proposed by McPeek will be implemented only if a specially selected, finely tuned parameter is used, and, therefore, “it is therefore not a robust mechanism”. It turns out that, according to the authors of the comments, the severity of the conditions of the model used excludes the reliability of the detected mechanisms, since the result of competition can change with changing environmental conditions. But this logic makes a thorough study of individual mechanisms unproductive. Each individual mechanism of competitive coexistence is implemented under certain conditions, and under these conditions it is reliable. References to the fact that in nature there may be a change in circumstances affecting the coexistence of competing species cannot serve as arguments against the reliability of the existence of certain mechanisms of competitive coexistence. Nature is not a swindling gamer playing in “pea and thimbles”, deceitfully changing the pea position. It is not the rules of the game that change, but the combinations of individual provisions of these rules (that is, combinations of mechanisms). Each individual mechanism can be identified, and its specific relationship with other mechanisms can also be identified. Only accurate and rigorous models can reveal rigorous and accurate environmental mechanisms.
4. And the fourth source of the difficulties is the high degree of uncertainty of the concept of “niche” (Haefner 1980, Leibold 1995, McInerny and Etienne 2012a, b, c, Olson et al. 2019). In a general sense, the term “niche” refers to the entire set of ecological characteristics of a species, including the characteristics of habitat (resources), characteristics of the resource needs, and functional characteristics of the adaptive abilities of the species. For accurate studies of the mechanisms of competitive coexistence in mathematical models, it is necessary to clearly separate functional traits of the species and their resource habitat.

These four sources of difficulties in understanding the mechanisms for implementing the principle of competitive exclusion served as the causes of the long period of uncertainty regarding the biodiversity paradox. Our paper presents the results of idealized model experiments, in which we were able to exclude at the basic level all four sources of the long uncertainty regarding validity of these two hypotheses.

The following conditions have been used in our idealized model experiments by which we try to exclude at the basic level all four sources of the uncertainty regarding the validity of the two hypotheses under consideration:

1. free micro-habitat is the only resource consumed by each individual of competing species during its life and the only resource for which interspecific competition occurs;
2. all free micro-habitats are identical;
3. each and everyone individuals of competing species had identical resource needs;
4. the only variable parameter of our model is the value of individual competitiveness, which is defined as the relative ability of an individual of a species to reproduce into a free micro-habitat in a conflict of interest.

### 1.2 Individual-based mechanisms of coexistence of complete competitors

We propose and investigate a white-box model of ecosystem dynamics supplemented by a probabilistic determination of the competitiveness of competing individuals without cooperative effects (Model 1) and its extension Model 2 with cooperative effects based on the numerical superiority of individuals of the species, locally competing for a given microhabitat. A feature of this model approach is its multilevel character. Along with colonization of free ecosystem starting from single individuals, we investigate cases when half of the habitat was probabilistically populated by equal numbers of individuals of both species at initial iteration. These models of ecosystem dynamics and resource competition between two species are mechanistic because they are individual-based and takes into account local interactions between individuals. In the case when species difference is maximal the model is completely deterministic. In this case the model is of the completely white-box type. In other cases, supplemented by a probabilistic determination of the competitiveness the models are of the grey-box type. Mathematical models of complex systems may be of three types in accordance with the degree of their mechanicalness – black-box, grey-box and white-box (Kalmykov and Kalmykov 2015c, Kalmykov and Kalmykov 2016).

Earlier we found two deterministic individual-based mechanisms of coexistence of complete competitors (Kalmykov and Kalmykov 2013, 2015b). *The first coexistence mechanism* is based on free resource gaps (free sites) which help to eliminate direct conflicts of interest between competing species, and as result colliding population waves of different competing species interpenetrate through each other like soliton waves in physical systems (Kalmykov and Kalmykov 2013). A possible mechanism of appearance of such gaps is moderate reproduction which was modelled through a hexagonal rosette-like cellular automata neighborhood. *The second coexistence mechanism* is based on timely recovery of the limiting resource, its spatio-temporal allocation between competitors and limitations of the habitat size (Kalmykov and Kalmykov 2015b). This mechanism allows complete competitors coexist in spite of their aggressive propagation without gaps in population waves. However, these mechanisms of indefinite coexistence are limited by the conditions of the *moderate propagation* and the particular *initial location of individuals on the lattice*. The principle must always be right and if there are exceptions, then either the principle is not true, or its formulation is not correct. The revealed mechanisms of competitive coexistence violate the known formulations of the competitive exclusion principle and, as result, we have reformulated the principle (Kalmykov and Kalmykov 2013).

### 1.3 A hypothesis

Previously we proved that complete competitors may coexist on one limiting resource regardless their 100% difference in competitiveness (Kalmykov and Kalmykov 2013, 2015b). However, these results are dependent on the moderate propagation of the competitors and on the particular initial location of individuals of the competing species in the ecosystem. Here we have demonstrated that reducing the difference in competitiveness may help to overcome this dependence. We study a hypothesis *that two completely competing species may coexist indefinitely with smaller than 100% difference in competitiveness regardless of their aggressive propagation and of random initial location of individuals of competing species in their ecosystem*.

## 2 METHODS

### 2.1 Methodology

A number of problems in theoretical ecology were caused by difficulties in mathematical modeling of complex systems. Such difficulties include the opacity (“phenomenological”) of mathematical modeling methods (Tilman 1987, Kalmykov and Kalmykov 2016), the complexity of modeling local cause-and-effect dependencies, the problems of integral multi-part and multi-level modeling of complex systems, the problems of modeling the behavior of multi-agent systems in which each individual agent makes its own decisions, the problems of individual-based modeling competition for ecosystem resources. Elements of arbitrariness in the interpretation of the black box models generated contradictions and paradoxes including the problem being solved here. In this paper, we have developed a method of transparent artificial intelligence to investigate mechanisms for implementing the competitive exclusion principle. This method allows us to implement automatic deductive inference based on a set of axioms of our theory of interspecific competition in the ecosystem. The basic definitions of our axiomatic theory are the concepts of micro-ecosystem and competitivity. Space is modeled by the lattice of a cellular automaton, whose cells correspond to micro-ecosystems, and time is modeled by successive iterations of the simultaneous change of States of all micro-ecosystems.

### 2.2 Basic definitions of ecosystem theory

*Ecosystem* is the unity of living organisms and their habitats interacting as a whole. The lattice of the cellular automata simulates an ecosystem.

*Micro-ecosystem* is the unity of an individual and its’ micro-habitat. This is an elementary basic object of the theory. In order to correctly model spontaneous changes in the object’s states, it is considered as a closed autonomous self-moving system at each time when a decision is made to change the state. A single lattice site (cell) simulates a one microecosystem.

*Meso-ecosystem* is the unity of a meso-habitat and individuals which inhabit in it. This basic object links changes in the states of micro-ecosystems into a single whole at the macro level by means of their local interactions. Meso-ecosystem modeled by the cellular automata neighborhood.

*Macro-ecosystem* is the unity of a macro-habitat and individuals which inhabit in it. It modeled by the cellular automata lattice.

*Habitat* is the intrinsic environment (the specific natural home) where a part of a species is able to survive.

*Micro-habitat* is the intrinsic environment where a particular individual inhabits. A micro-habitat is a totality of all environmental conditions and resources which are necessary for individual’s life (e.g. for a grass unit) and place where regeneration of the resources is possible. A free micro-habitat represents a place which can be occupied by an individual autonomous agent (e.g. by an individual of a species) and contains the necessary resources for its individual life. A microhabitat inhabited by an individual correspond to the microecosystem.

*Meso-habitat* is the intrinsic environment where a particular individual of a species is able to propagate in one generation. Here, in the cellular automaton models a mini-habitat is a cell together with its neighboring cells but without an individual itself. Mesohabitat with individuals forms a meso-ecosystem.

*Macro-habitat* is the total modeling environment where individuals of the species may inhabit and propagate potentially and actually. Here, in the cellular automata model, macro-habitat is the entire field of the cellular automata without competing individuals. Macro-habitat contains a complete set of all micro-habitats of the modeling ecosystem. Macro-habitat with individuals living in it forms macro-ecosystem.

*Individual* is one organism of a species (an autonomous agent in the most general case).

*Competition for resources* is a type of rivalry relationship that occurs when at least two parties seek to consume the same limiting resource and when the benefit received by one side of the conflict of interest is the loss of the other. In our model, competition for resources occurs between individuals of competing species in a direct conflict of interest for possibility of reproduction into a free micro-habitat.

*Competitiveness of an individual* is defined as the relative ability of the individual of a species to reproduce into a free micro-habitat in a direct conflict of interest with an individual of the competing species. The relative competitiveness of individuals is the result of differences in their adaptive functional characteristics, and this is the only interspecific difference between individuals in our model.

*Fecundity* is the potential for reproduction - the potential ability to produce offsprings of an individual or population. A cellular automata neighborhood determines the number of possible offsprings. Fecundity rate is a potential per capita offsprings’ number on a next iteration.

*Fertility* is the actual production of offsprings - the actual reproductive rate of an individual or population. Fertility rate is an actual per capita offsprings’ number on a next iteration.

### 2.3 Axioms that specify interactions between objects defined by basic definitions

1. *A space* is modelled by a lattice with microecosystems at its nodes.
2. *A time* is modelled by sequential iterations of changes of states of all microecosystems.
3. *List of micro-ecosystem states*:
- ‘Free’ – may be occupied only by one immobile individual of any species;
- ‘Occupied’ – the occupied micro-ecosystem goes into the regeneration state after an individual’s death;
- ‘Regeneration’ – restoring conditions and resources of a microecosystem after an individuals’s death and recycling of a dead individual of a species. It cannot be directly occupied. Micro-ecosystem in the regeneration state becomes free or may be occupied immediately after finishing of the regeneration state
4. *Neighborhood of local interactions* of micro-ecosystems with each other;
5. *Rules for changing micro-ecosystem states depending on their local neighborhood.*

### 2.4 Automatic deductive inference

Automatic deductive inference is carried out through the formalism of logical cellular automata. Cellular automata are mathematical idealizations of physical systems in which space and time are discrete, and physical quantities take on a finite set of discrete values (Wolfram 1983). The formalism of logical cellular automata includes:

1. *A regular uniform lattice of cells* (micro-objects of the model);
2. *A finite set of possible states of a cell*;
3. *Neighborhood*, consisting of a cell and its’ certain neighboring cells. This set of cells have an influence to the each next-state transition of a cell. Neighborhoods of all cells are the same;
4. *The next-state transition function* – the rules for iterative changes of the states of each and every cell is applied to the whole lattice simultaneously. These rules taking into account the state of the neighborhood of each cell. All cells have the same rules for updating. The iterations of the cell’s states model a time scale. A logical cause-and-effect structure governing the state-transition of a cell is based on *if-then* statement.

*Each cell of a cellular automata is a finite automaton* (synonyms – a finite-state machine, a finite-state automaton or simply a state machine), and the cellular automaton as a whole is a *polyautomaton* which co-organizes the behavior of the finite automata included in it. Deductive inference is implemented as hyper-logics because cellular-automatic logic is simultaneously performed as logical inference operations with respect to each and every micro-object of the modeled system at each iteration of the model evolution. The presented method of automatic axiomatic deductive inference is a promising method of artificial intelligence. This is the realization of the old dream of symbolic artificial intelligence “GOFAI” - Good Old-Fashioned Artificial Intelligence (Haugeland 1985). Our artificial intelligence method overcomes the opacity problems that are characteristic of solutions in artificial neural networks. Unlike indirect solutions to overcome the opacity of artificial intelligence methods in the framework of the “Explained Artificial Intelligence (XAI)” project (Adadi and Berrada 2018), our method allows us to directly implement ecologically inspired transparent artificial intelligence in the framework of the white-box type model. We view our version of transparent artificial intelligence as the most advanced version of explainable artificial intelligence (XAI) as we directly and in all possible details obtain understanding of mechanisms underlying the system under study.

### 2.5 A biological prototype of the model

A vegetative propagation of rhizomatous lawn grasses is the biological prototype of our models. *Festuca rubra trichophylla* (Slender creeping red fescue) is the prototype of aggressive vegetative propagation and *Poa pratensis L.* and *Festuca rubra L. ssp. Rubra* are the prototypes of moderate vegetative propagation. Behind vegetative propagation, we would like to see a more general case of reproduction. Vegetative reproduction is interesting to us not in itself, but only as a way to exclude the emergence of the genetic diversity of individuals of a species. In addition, we were attracted by the ability of rhizomatous lawn grasses to the spatially ordered arrangement of descendants in the form of the two-dimensional lattice structures. One individual corresponds to one tiller. A tiller is a minimal relatively autonomic grass shoot that sprouts from the base of grass and which is able to propagate. Rhizomes are horizontal creeping underground shoots using which plants vegetatively (asexually) propagate. Unlike a root, rhizomes have buds, scaly leaves and nutrient reserves for semiautonomous development. A tiller with roots and leaves develops from a bud on the end of the rhizome. One tiller may have maximum six rhizomes in the model. Six rhizomes per tiller correspond to aggressive vegetative propagation. A populated micro-habitat goes into the regeneration state after a tiller’s death. The regeneration state of a site corresponds to the regeneration of micro-habitat’s resources including recycling of a dead individual.

### 2.6 A transparent individual-based modeling

For quantitative verification of these hypotheses, we use here a transparent individual-based modeling approach based on the cellular automata logic. The transparency means not only that individuals are present in the model, but also that we can control the dependence of each individual’s behavior on its local conditions. This allows us to study mechanisms of interspecific competition in a quantitative and the most complete way, taking into account the relative competitiveness of each and everyone individual. The general methodology of the transparent modeling of complex systems can be found in our early works (Kalmykov and Kalmykov 2013, 2015b, c, a, Kalmykov and Kalmykov 2016, Kalmykov et al. 2017). An elementary object of our models is a microecosystem. This is the basic ideal “self-moving” object. The resource closure of a microecosystem allows to consider it as an autonomous object. Autonomous dynamics of this object is characterized by a set of states, and the diagram of their sequential change in accordance with the physical principles of the extreme. Spontaneous dynamics of this autonomous object is modeled as transitions between the states of a lattice site (cell). Transitions between the states are accompanied by use of free energy. The free energy is used to ensure the livelihoods of individuals (the high-grade energy of solar radiation) and to maintain processes of the limiting resources recovery during a regeneration state of a microhabitat (energy of solar radiation and energy of organic matter of dead individuals). An individual of the dominant species has greater ecological competitiveness and wins in a direct conflict for resources for propagating. It has the most effective ability of capturing and using free energy and other limiting resources. Following (Tansley 1935), we consider individuals with their “special environment, with which they form one physical system”. A microhabitat is the intrinsic part of environmental resources of any one individual and it contains all necessary resources for its life. It is an intrinsic part of all environmental resources and conditions per one individual. The high-grade (“fresh”) energy flow of solar radiation provides an individual’s life in a microhabitat. The flow of lowered-grade (“used-up”) energy which is produced during an individual’s life goes into the cold space and also provides the conditions for an individual’s life in a microhabitat. A living organism is not able to function without the heat output to a “refrigerator”. Ultimately, the cold space performs a role of such refrigerator. A working cycle of a microecosystem’s processes includes occupation of a microhabitat by an individual, life of the individual and restoration of used resources after an individual’s death. A microecosystem includes a single microhabitat and is able to provide a single accommodation of one individual, but it cannot provide its propagation. This is due to the fact that after an individual’s death, its microhabitat goes into a regeneration (refractory) state, and is not suitable for immediate occupation. A minimal ecosystem, providing a possibility of vegetative propagation of an individual, must consist of two microhabitats. The vegetative propagation is carried out due to resources of an individual’s microhabitat by placing a vegetative primordium of a descendant in a free microhabitat of an individual’s neighborhood. A plant ecosystem we consider as “a working mechanism” which “maintains and regenerates itself” (Watt 1947). To simulate vegetative propagation, we use the axiomatics of excitable medium (Wiener and Rosenblueth 1946) and the concept of an individual’s intrinsic microecosystem (Kalmykov and Kalmykov 2015b). In accordance with this axiomatics, the three states of a medium - the rest, excitation, and refractoriness successively replace each other. The rest state corresponds to the free state of the microecosystem, the excitation state corresponds to the life of an individual in the microecosystem, and the refractory state corresponds to the state of regeneration of the microecosystem after the death of an individual. After an individual’s death, its microecosystem goes into a regeneration (refractory) state, which is not suitable for immediate occupation. In our model vegetative propagation is carried out due to resources of an individual’s microecosystem by placing its vegetative primordium of a descendant in a free micro-habitat of an individual’s neighborhood. A microecosystem includes a single micro-habitat and is able to provide accommodation of one individual. Taking into account the refractory state of micro-habitat expands possibilities of individual-based modeling of ecosystem dynamics because most ecological models still do not take into account regeneration of resources in plant communities. This problem was first discussed in general terms by Watt in 1947 (Watt 1947, Grubb 1977).

Cellular automata are ‘bottom up models’ that generate global behavior from local rules (Silvertown et al. 1992). In this study, we have carefully developed the rules for these local dependencies in order to obtain unambiguous conclusions about the validity of the hypotheses under consideration. Due to the ‘bottom up’ inference, we were able to ensure that our model mechanically constructs an ecosystem at each iteration from the local conditions of each individual micro-environment and of each and every individual to the ecosystem as a whole. The rules of cellular automata are implemented at three levels of ecosystem organisation. It is this multi-level modeling that allows us to talk about the transparency of this method (Kalmykov and Kalmykov 2013, 2015b, c, a, Kalmykov and Kalmykov 2016, Kalmykov et al. 2017). A micro-level is modelled by a lattice site (micro-ecosystem). A meso-level of local interactions of a micro-ecosystem is modelled by the cellular automata neighborhood (meso-ecosystem). A macro-level (the entire ecosystem) is modelled by the entire lattice. A two-dimensional hexagonal lattice is closed to a torus by periodic boundary conditions in order to avoid boundary effects. We use the hexagonal lattice because it most naturally implements the principle of densest packing of micro-habitats. The hexagonal neighborhood allows to model a potentially aggressive vegetative propagation of plants when offsprings of an individual may occupy all nearest micro-habitats (Fig. 1a, Movies S1-S3).

**Figure 1.**
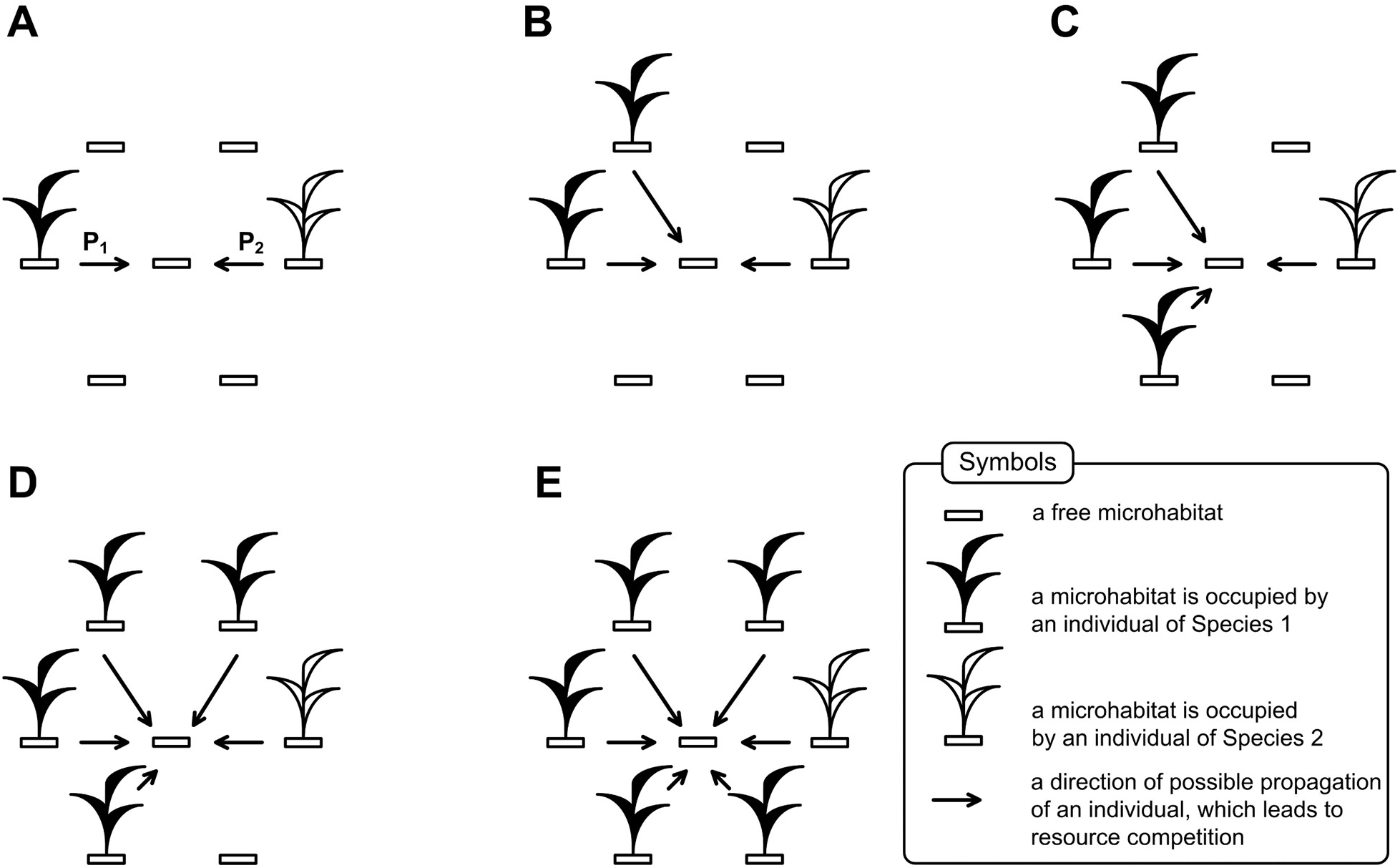
A schematic representation of interspecific competition for a free microhabitat in Model 1. An increase in the number of individuals of a particular species that can occupy a free microhabitat does not increase chances of this species to occupy it. Parameters P1 and P2 represent probabilities of occupation of the free microhabitat by an offspring of Species 1 and Species 2, respectively. The maximum number of individuals, competing for the single microhabitat equals six (E). Chances to win in a direct conflict of interest are independent from the number of competing individuals – parameters P_1_ and P_2_ are constant for all cases (A-E).

### 2.7 The cellular automata Model 1 of resource competition without cooperative effects

In order to test our hypothesis we use our individual-based cellular automata model of interspecific competition in conditions that exclude the implementation of the multiple known mechanisms of competitive interspecies coexistence (Hutchinson 1961, Hastings 1980, Palmer 1994, Chesson 2000, Wellborn 2002, Dollhopf et al. 2003, Nowak 2006, Bennett and Bever 2009) (Figs. 1, 2):

- A habitat has a limited size, it is homogeneous (micro-habitats are identical) and stable (i.e. environmental conditions are constant, – there are no any climate or abiotic resources fluctuations and as a consequence relative competitiveness of competing species remain unchanged;
- Immigration, emigration, predation, herbivory, parasitism, infectious diseases and other disturbances are absent;
- Competing species are per capita identical and constant in ontogeny, in fecundity (potential fertility) rates, in regeneration features of habitats and environmental requirements.
- Reproduction of the competing species occurs only vegetatively and the species are genetically homogeneous and stable (from generation to generation there is no change of heredity);
- We model the species which are trophically identical consumers and differ only in relative competitiveness of their individuals;
- There are no any trade-offs, mutualism and cooperative interactions between individuals of the competing species;
- Offsprings of an individual potentially may vegetatively occupy all of the nearest micro-habitats. Such propagation is aggressive and it is modelled by the hexagonal neighborhood.

**Figure 2.**
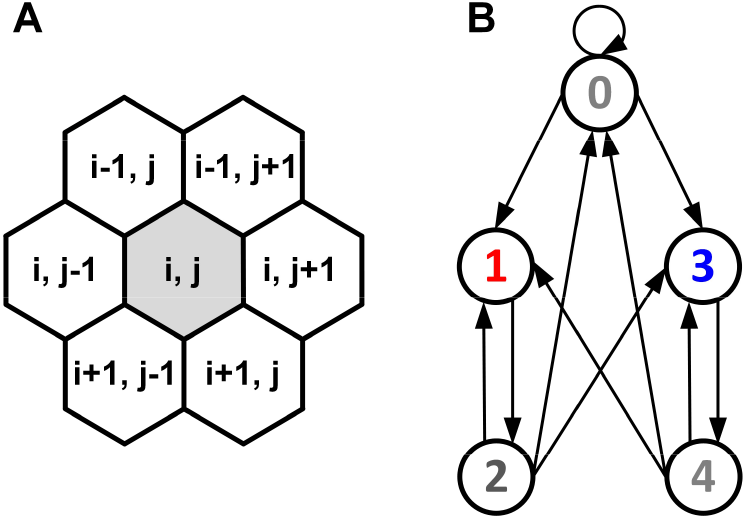
The hexagonal neighborhood and the diagram of transitions between the states of a site. (A) The hexagonal neighborhood. The central site of the neighborhood is defined by the array element with index (i, j), where i and j are integer numbers. Neighboring sites of the central site are defined by the array elements with indexes (i - 1, j), (i - 1, j + 1), (i, j + 1), (i + 1, j), (i + 1, j - 1), (i, j - 1). Possible local interactions between competing individuals are defined by this neighborhood. (B) A directed graph of logical probabilistic transitions between the states of a lattice site. Each state and transition between states have a specific interpretation.

Additionally, multiple diversity of ecological conditions in our model experiments is extremely minimized to give clear conclusions – all individuals of both competing species are considered as identical consumers, competing, all other things being equal, for a single limiting resource - a free micro-habitat. Prior to the initial settlement of individuals, the habitat is homogeneous and consists of identical free micro-habitats. The main variable parameter is a value of the species’ competitiveness which is defined as a relative ability of an individual of the species to use a single free micro-habitat for propagating in a local conflict of interests. The only difference between competing species is competitiveness of their individuals, which in our model is supposed to be a consequence of differences in their adaptive functional traits.

Our goal is a theoretical search for cases of coexistence of two complete competitors under conditions which are the most unfavorable for coexistence. One of conditions in verifying the principle was the exclusion of cooperative effects. It is known that cooperation can be a mechanism for ensuring the coexistence of competitors with different competitiveness (Lampert and Tlusty 2011). To exclude the contribution of cooperative effects in Model 1, we have established that the probability of winning does not depend on how many individuals of a species compete for this microhabitat (Fig. 1).

Trade-offs are one of the mechanisms of competitive coexistence (Amarasekare 2004) – so their absence is necessary to ensure strict conditions for verifying the validity of the classical formulations of the principle. We define competitiveness on an individually-oriented basis as a relative ability of an individual of a species for using a single free microhabitat for reproduction in the local interspecific competition. We assume that priority effects and some particular species traits may be mechanisms of the modeled difference in competitiveness. A priority effect may be the impact that a particular species can have on community development due to prior arrival at a site, for example, due to inhibitory priority effects when a species that arrives first at a site negatively impacts a species that arrives later by reducing the availability of space or resources (Fukami et al. 2005). A distinguishing feature of the ability to capture a resource, may also be a consequence of a specific functional eco-physiological traits - such as rate and strength of growth (Violle et al. 2007).

In our experiments, the maximum number of individuals of both species locally competing for one free habitat is six, the species are genetically homogeneous and the reproduction occurs asexually (vegetatively). Usually ecologists define competitiveness as the average reproductive success of a population of this species since classical mathematical methods do not allow to evaluate the reproductive success of a species on an individually-oriented basis. In accordance with the neighborhood of the cellular automata, the fecundity of each individual of a given species at a given iteration can vary from zero to six. This value is determined by the presence of free sites in the neighborhood of an individual, and also by presence of the competitor individuals claiming for these free sites. If there are free microhabitats in the neighborhood of this individual, and individuals of a competing species do not claim them, then at the next iteration the descendants of this individual occupy them. If the conflict of interest arises, then in Model 1 for each individual free site the outcome of this conflict is determined probabilistically in accordance with constant competitiveness parameters P_1_ and P_2_ of each species without considering how many individuals of each species compete for the microhabitat.

A vegetative propagation of rhizomatous lawn grasses is the biological prototype of our models. Behind vegetative propagation, we would like to see a more general case of reproduction. Vegetative reproduction is interesting to us not in itself, but only as a way to exclude the emergence of the genetic diversity of individuals of a species. In addition, we were attracted by the ability of rhizomatous lawn grasses to the spatially ordered arrangement of descendants in the form of lattice type structures. One individual corresponds to one tiller. A tiller is a minimal relatively autonomic grass shoot that sprouts from the base of grass and which is able to propagate. Rhizomes are horizontal creeping underground shoots using which plants vegetatively (asexually) propagate. Unlike a root, rhizomes have buds, scaly leaves and nutrient reserves for semiautonomous development. A tiller with roots and leaves develops from a bud on the end of the rhizome. One tiller may have maximum three or six rhizomes in the model. Three rhizomes per tiller correspond to moderate propagation only in a half of the nearest microhabitats and six rhizomes per tiller correspond to aggressive vegetative propagation (Fig. 2A). *Festuca rubra trichophylla* (Slender creeping red fescue) is the prototype of aggressive vegetative propagation and *Poa pratensis L.* and *Festuca rubra L. ssp. Rubra* are the prototypes of moderate vegetative propagation. A populated microhabitat goes into the regeneration state after an individual’s death. The regeneration state of a site corresponds to the regeneration of microhabitat’s resources including recycling of a dead individual. All individuals are identical.

In Model 1 an individual of the Species 1 has a probability P_1_ to win in a direct conflict for resources, and an individual of the Species 2 has a probability P_2_ to win in the same conflict (Figs. 1, 2B). Parameter P1 reflects competitiveness of the Species 1 and parameter P_2_ reflects competitiveness of the Species 2. In all cases here, a microhabitat under the conflict of interest will always be occupied by an individual of one of the species, so P_1_ + P_2_ = 1. By changing competitiveness parameter P_1_ we looked for a maximum value of competitiveness difference when competing species can coexist. The competitiveness difference is determined as P_1_ - P_2_. Starting from the deterministic case of competitive exclusion when P_1_ = 1 we investigated most revealing cases of competitive coexistence with an increment of competitiveness difference which equals 0.01.

Here we extend our logical cellular automata model of ecosystem with two competing species (Kalmykov and Kalmykov 2013, 2015b) supplementing them by a probabilistic determination of the competitiveness of competing individuals (Model 1). Here we introduce probabilistic rules of competition to investigate the influence of competitiveness differences on the species coexistence. The model is based on (i) the formalism of excitable medium and (ii) on the concept of an individual’s intrinsic microecosystem (Kalmykov and Kalmykov 2015b). The entire cellular automaton simulates a whole ecosystem which autonomously maintains and regenerates itself. A two-dimensional hexagonal lattice is closed to a torus by periodic boundary conditions in order to avoid boundary effects. We use the hexagonal lattice because it most naturally implements the principle of densest packing of microhabitats. The hexagonal neighborhood allows to model a potentially aggressive vegetative propagation of plants when offsprings of an individual may occupy all nearest microhabitats (Fig. 2A). Each site of the lattice simulates a microhabitat. A microhabitat contains resources for existence of a one individual of any species. A microecosystem is a microhabitat with an individual living in it. A microhabitat is the intrinsic part of environmental resources of one individual and it contains all necessary resources for its autonomous life. An individual can occupy a one microhabitat only. A life cycle of an individual lasts a single iteration of the automaton. All states of all sites have the same duration. Every individual of all species consumes identical quantity of identical resources by identical way, i.e. they are identical per capita consumers. Such species are complete competitors. Individuals are immobile in lattice sites and populations waves propagate due to reproduction of individuals (Supplemental Videos S1-S3). The closest biological analogue is vegetative reproduction of rhizomatous lawn grass (Fig. 1).

A neighborhood consists of a site and its intrinsically defined neighbor sites (Fig. 2A). All sites have the same rules for updating. А neighborhood simulates an individual’s mini-habitat and determines the number of possible offsprings (fecundity) of an individual.

Here is a description of the states of a lattice site (Fig. 2B). Each site may be in one of the four states:
0—a free microhabitat which can be occupied by a single individual of any species;
1—a microhabitat is occupied by a living individual of the Species 1;
2—a regeneration state of a microhabitat after the death of an individual of the Species 1;
3—a microhabitat is occupied by a living individual of the Species 2;
4—a regeneration state of a microhabitat after the death of an individual of the Species 2.

Rules of transitions between the states of a site of the two-species competition model (Fig. 2):

0 → 0, A microhabitat remains free if there is no one living individual in the neighborhood;

0 → 1, If in the cellular automata neighborhood are individuals of both competing species, then the probability of this transition is defined by the parameter P_1_. If in the cellular automata neighborhood is at least one individual of the Species 1 and there is no one individual of the Species 2, then this transition is always implemented;

0 → 3, If in the cellular automata neighborhood are individuals of both competing species, then the probability of this transition is defined by the parameter P_2_. If in the cellular automata neighborhood is at least one individual of the Species 2 and there is no one individual of the Species 1, then this transition is always implemented;

1 → 2, After the death of an individual of the Species 1 its microhabitat goes into the regeneration state;

2 → 0, After the regeneration state a microhabitat will be free if there is no one living individual in the neighborhood;

2 → 1, If in the cellular automata neighborhood are individuals of both competing species, then the probability of this transition is defined by the parameter P1. If in the cellular automata neighborhood is at least one individual of the Species 1 and there is no one individual of the Species 2, then this transition is always implemented;

2 → 3, If in the cellular automata neighborhood are individuals of both competing species, then the probability of this transition is defined by the parameter P2. If in the cellular automata neighborhood is at least one individual of the Species 2 and there is no one individual of the Species 1, then this transition is always implemented;

3 → 4, After the death of an individual of the Species 2 its microhabitat goes into the regeneration state;

4 → 0, After the regeneration state a microhabitat will be free if there is no one living individual in its neighborhood;

4 → 1, If in the cellular automata neighborhood are individuals of both competing species, then the probability of this transition is defined by the parameter P1. If in the cellular automata neighborhood is at least one individual of the Species 1 and there is no one individual of the Species 2, then this transition is always implemented;

4 → 3, If in the cellular automata neighborhood are individuals of both competing species, then the probability of this transition is defined by the parameter P2. If in the cellular automata neighborhood is at least one individual of the Species 2 and there is no one individual of the Species 1, then this transition is always implemented.

These logical statements include probabilistic parameters P_1_ and P_2_ which allow to investigate a role of competitiveness differences in coexistence of two aggressively propagating and competing species.

The cellular automata rules are realized on the three levels of organization of the complex system. A micro-level is modelled by a lattice site (micro-ecosystem). A mini-level of local interactions of a micro-ecosystem is modelled by the cellular automata neighborhood (mini-ecosystem). A macro-level (the entire ecosystem) is modelled by the entire lattice.

Different initial conditions may lead to formation of different spatio-temporal patterns and, as a result, to different dynamics of the system. We have investigated two situations on initial iteration – (i) when colonization of free habitat started from a single individual of each species (Fig. 3A) and (ii) when 25% of the territory was probabilistically populated by individuals of the Species 1, 25% of the territory was probabilistically populated by individuals of the Species 2 and 50% of the territory remained free (Fig. 3B). Here in Fig. 3 we show examples of initial patterns when the lattice consists of 50×50 sites.

**Figure 3.**
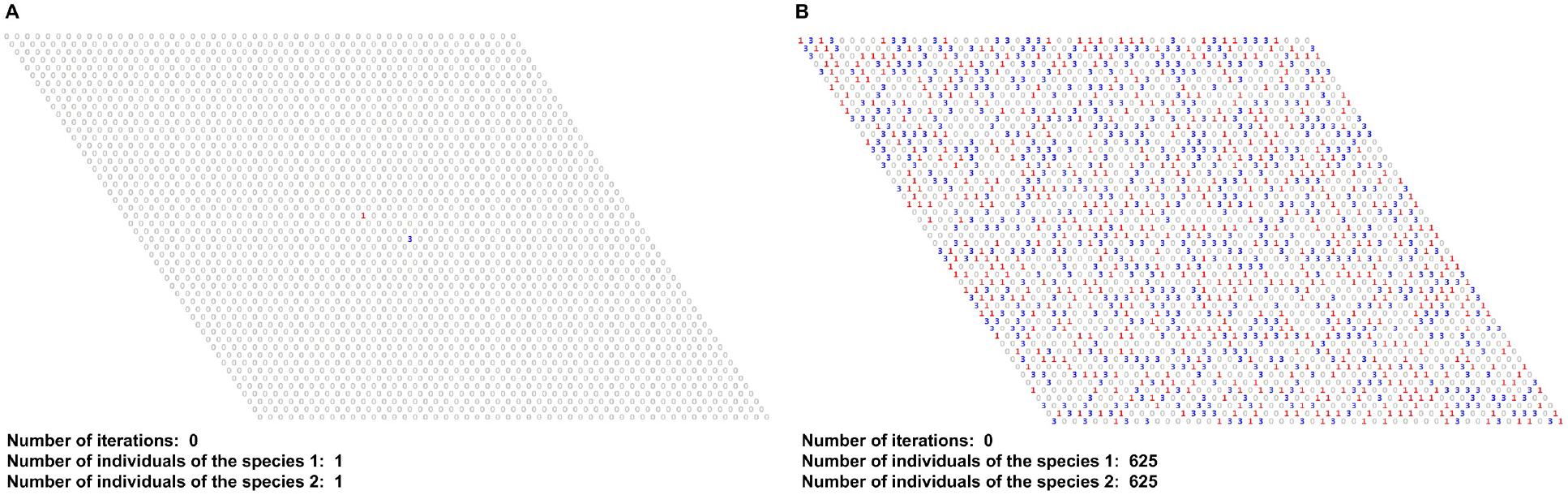
Examples of initial patterns. Cellular automata lattices consist of 50×50 sites and closed on the torus. (A) The cellular automata field was populated by single individuals of two competing species on initial iteration. (B) The field was probabilistically populated by 625 individuals of each of two competing species on initial iteration, i.e. 25% of the territory is occupied by individuals of the Species 1, 25% of the territory is occupied by individuals of the Species 2 and 50% of the territory is free.

In order to show that competing species coexist stably regardless of the initial placement of individuals in the habitat, we use a Monte Carlo simulation. The Monte Carlo simulation consists of 200 trials, when each trial lasts 10000 iterations.

We present two source codes of our Model 1 – completely competing species with competitiveness inequality written in C++ programming language (see Appendix S1).

#### Source code 1

Coexistence of complete competitors with 10% difference in competitiveness without cooperative dependence of competitiveness on the number of locally competing individuals. The initial pattern of the habitat is populated by single individuals of two competing species.

#### Source code 2

Coexistence of complete competitors with 10% difference in competitiveness without cooperative dependence of competitiveness on the number of locally competing individuals. The initial pattern of the habitat is randomly populated by individuals - 25% by the Species 1, 25% by the Species 2 and 50% of the habitat remains free.

The source code of the John Conway’s Game of Life (Naumov and Shalyto 2003a, Naumov and Shalyto 2003b) was partially used as a starting point for designing of the programs.

### 2.8 The cellular automata Model 2 of resource competition with cooperative effects

The main results of the study were found in conditions in which we excluded the participation of all known mechanisms of competitive coexistence of trophically similar species. This abstraction was mainly focused on checking the validity of the classical formulations of the competitive exclusion principle. For additional verification of our results obtained on the assumption that the competitiveness of the species does not depend on the number of individuals locally competing for a given microhabitat, we have investigated cases with a cooperative dependence. Such verification was made in order to make our abstract models more realistic. It was interesting to investigate possible cases of competitive coexistence under conditions with cooperative effects in our models. To do this we extended Model 1 to Model 2 supplementing Model 1 by cooperative effects based on the numerical superiority of individuals of the species, locally competing for a given microhabitat. In these additional experiments the number of individuals of one species locally competing with another species affects the chances of this species to propagate. Cooperative effects are based on the numerical superiority of individuals of this species, locally competing for a given microhabitat. Competitiveness of the species increases in proportion to its numerical superiority. We proceeded from the assumption that an increase in the competitiveness of the species at its maximum numerical superiority should ensure the local victory of its individual, all other things being equal.

The maximum possible numerical superiority is the ratio of 5 individuals of one species to 1 of another (see Fig. 1E) – the difference in numbers of individuals in this case is 4.

We investigated influence of cooperative effects on interspecific competition in two cases, differing in initial patterns (see 3.3.6 in Results):
1. Initial pattern was populated by two single individuals of competitors. Without cooperative effects we have found that the extinction of the recessive species begins with 32% difference in competitiveness. Therefore, investigating cooperative effects, we have defined the competitiveness increment with an increase in the number of individuals by 1 unit as 32%:4=8%.
2. Initial pattern was randomly populated by equal number of individuals of competitors. Without cooperative effects we have found that the extinction of the recessive species begins with 23% difference in competitiveness. Therefore, we have defined the value of the competitiveness increment with an increase in the number of individuals by 1 unit as 24%:4 = 6%.

In each of these two cases, three variants of the starting basic differences of the competing species in competitiveness were studied: A - no differences, B - 10% difference and C - 20% competitiveness differences.

We present two source codes of the Model 2 – completely competing species with competitiveness inequality and with cooperative effects. They are written in C++ programming language (see Appendix S1).

#### Source code 3

Coexistence of complete competitors with the initial 10% difference in basic competitiveness and with dependence of the competitiveness on a number of locally competing individuals. The initial pattern of the habitat is populated by two single individuals of two competing species. The increase in the number of individuals of a species, locally competing for reproduction in a given free microhabitat by one unit, increases its competitiveness by 0.08 (by 8%).

#### Source code 4

Coexistence of complete competitors with the initial 10% difference in basic competitiveness and with dependence of the competitiveness on a number of locally competing individuals. The initial pattern of the habitat is randomly populated by individuals - 25% by the Species 1, 25% by the Species 2 and 50% of the habitat remains free. The increase in the number of individuals of a species, locally competing for reproduction in a given free microhabitat, by one unit, increases its competitiveness by 0.06 (by 6%).

## 3 RESULTS

### 3.1 Preliminary verification of the main hypothesis

Here we investigate the hypothesis that two completely competing species can coexist with smaller than 100% difference in competitiveness regardless of initial positioning of individuals in the habitat. We were changing the interconnected competitiveness parameters P_1_ and P_2_ to find a range of cases of coexistence of complete competitors. The most typical outcomes of competition obtained by varying the competitiveness parameter P1 are shown in Fig. 4. We investigated these outcomes of competition starting with one and the same initial positioning of individuals on the lattice. The most typical outcomes of competition are (i) the case of competitive exclusion (Fig. 4A and Supplemental Video S1) and the case of competitive coexistence of complete competitors with competitiveness inequality (Fig. 4B and Supplemental Video S2) and the case of competitive coexistence of complete competitors with equal competitiveness (Fig. 4C and Supplemental Video S3). Figure 4 demonstrates three cases of population dynamics in computer experiments presented in Supplemental Videos S1-S3, respectively.

**Figure 4.**
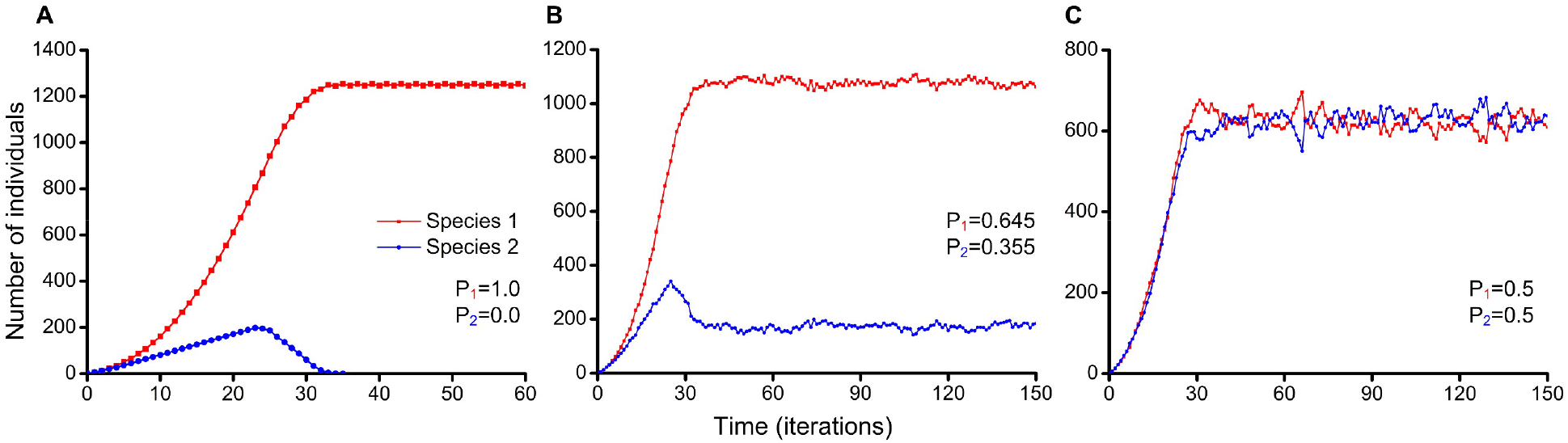
Population dynamics of two competing species. Colonization of free habitat which consists of 50×50 microhabitats started from a single individual of the Species 1 and a single individual of the Species 2. Parameter P_1_ reflects competitiveness of the Species 1 and parameter P_2_ reflects competitiveness of the Species 2. (A) The completely deterministic case of competitive exclusion when the Species 1 excludes the Species 2 (Supplemental Video S1). Species difference in competitiveness equals 100%. (B) The Species 1 and the Species 2 stably coexist despite the difference in competitiveness which equals 29% (Supplemental Video S2). (C) The Species 1 and the Species 2 are identical in all parameters and they stably coexist with close numbers of individuals (Supplemental Video S3). Species difference in competitiveness equals 0%.

### 3.2 Contribution of stochastic fluctuations in coexistence in small ecosystems

In order to check the influence of the lattice size in the contribution of stochastic fluctuations on results of competition, we investigated competition of two species with *the same competitiveness* in four small ecosystems with different lattice sizes (Fig. 5).

**Figure 5.**
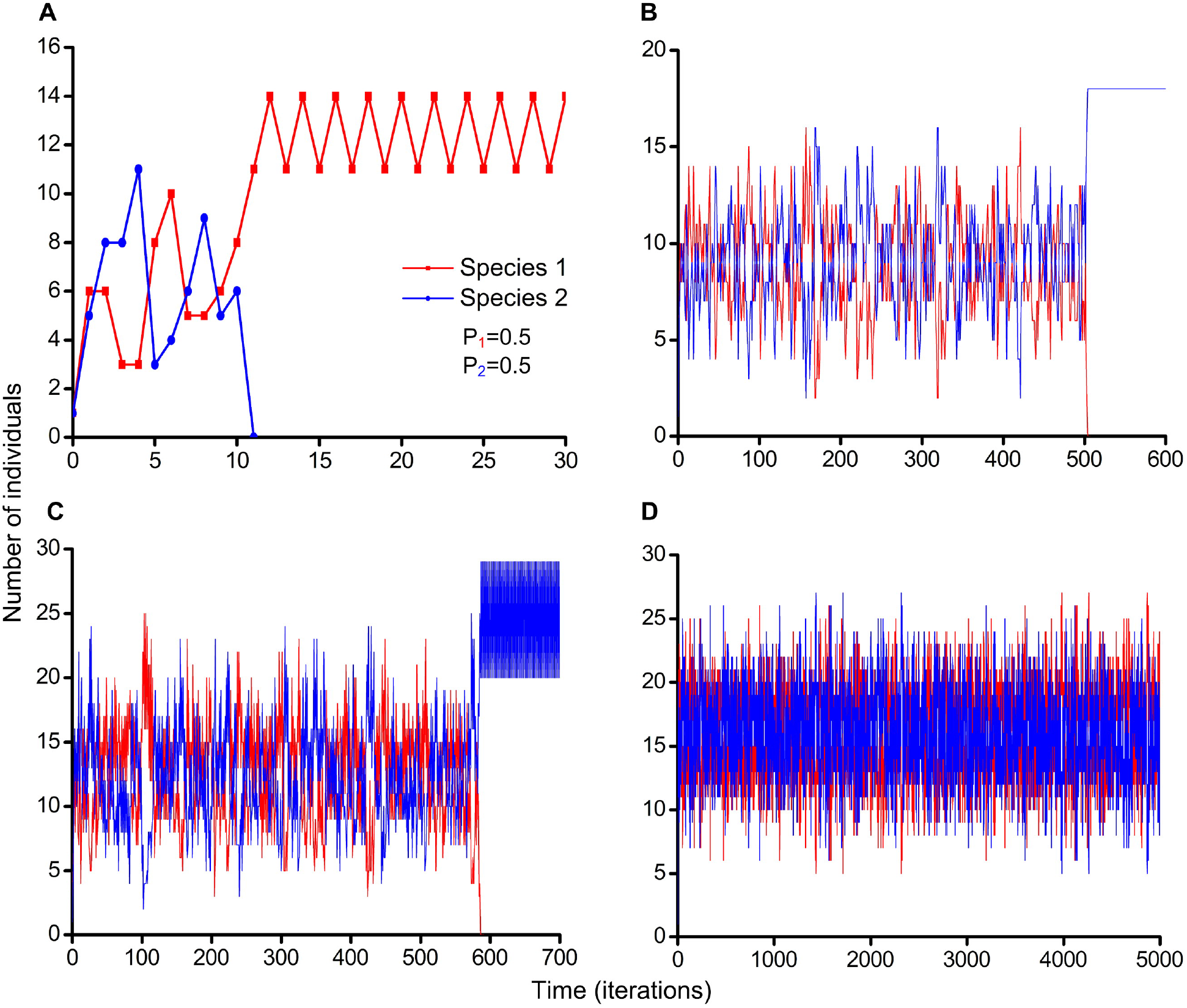
Influence of the habitat size in contribution of stochastic fluctuations on coexistence of two identical resource competitors. Colonization of free habitat started from single individuals of each species. Parameter P_1_ = 0.5 reflects competitiveness of the Species 1 and parameter P_2_ = 0.5 reflects competitiveness of the Species 2. Raw data is available on Figshare in Table S1 (https://doi.org/10.6084/m9.figshare.4903013.v1). (A) The Species 1 excludes the Species 2. The lattice consists of 5×5 sites. (B) The Species 2 excludes the Species 1. The lattice consists of 6×6 sites. (C) The Species 2 excludes the Species 1. The lattice consists of 7×7 sites. (D) The Species 1 and the Species 2 stably coexist. The lattice consists of 8×8 sites.

### 3.3 In-depth testing of the hypothesis with increased number of iterations and increased lattice sizes

#### 3.3.1 Monte Carlo simulations on the lattices 50×50 and 300×300 sites

Since with the increasing number of iterations and lattice sizes the revealed results become more objective, we repeated the studies presented in Fig. 4 with increased number of iterations from 150 up to 10000 and the lattice size from 50×50 to 300×300 sites (Figs. 6–9). These refinement experiments were carried out as Monte Carlo simulations with 200 trials in each case. The lattice was iteratively populated with probability 0.5% for each site before the criterion is reached – 625 individuals on the lattice 50×50 sites (Figs. 3B and 8) and 22500 individuals on the lattice 300×300 sites (Fig. 9) of each of two competing species. Our models differ from the Moran’s models (Moran 1958) in following points:
1. total number of individuals is not constant from iteration to iteration;
2. the lattice (habitat) is large enough to exclude the contribution of genetic drift;
3. the dynamics of each individual is based on local interactions between individuals.

**Figure 6.**
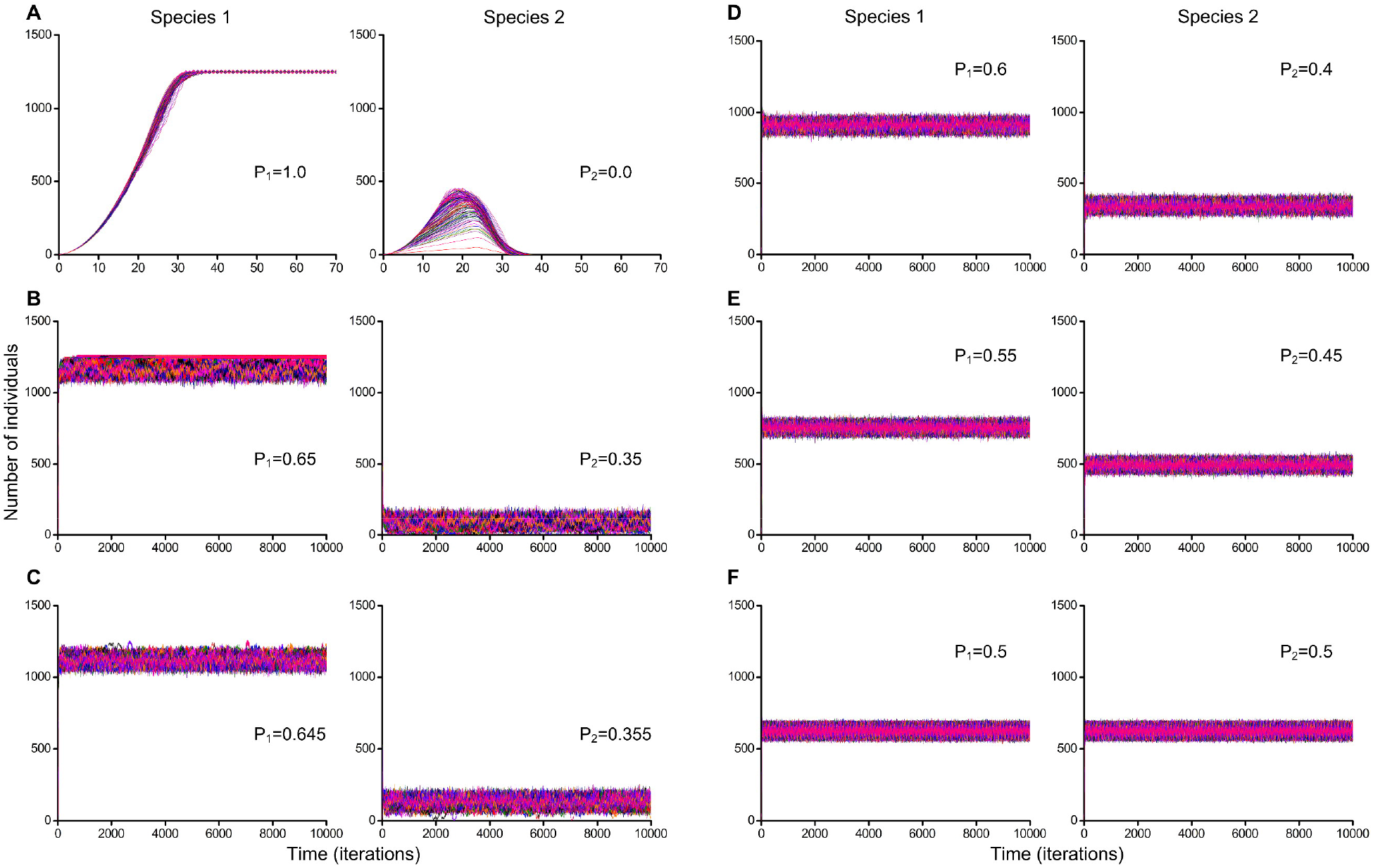
Monte Carlo simulations of two species competition with colonization of free habitat consisted of 50×50 sites. Colonization of free habitat started from single individuals of each species. The number of Monte Carlo simulations equals 200. Parameter P_1_ reflects competitiveness of Species 1 and parameter P_2_ reflects competitiveness of Species 2. Raw data is available on Figshare in Table S2 (https://doi.org/10.6084/m9.figshare.4902986.v1). (A) Competitive exclusion of the recessive Species 2 was observed in all cases. Species difference in competitiveness equals 100%. (B) There were 78 cases of competitive exclusion and 122 cases of competitive coexistence. Species difference in competitiveness equals 30%. (C-F) Competing species stably coexisted in all cases. Species differences in competitiveness equal 29% (C), 20% (D), 10% (E) and 0% (F).

Using the Monte Carlo method, we have investigated the influence of random initial positioning of individuals on the lattice (Figs. 6–9). In Figs. 6 and 7 every Monte Carlo simulation consisted of 200 repeated experiments with random initial positioning of a single individual of the Species 1 and a single individual of the Species 2 on the 50×50 lattice (Fig. 6) and on the 300×300 lattice (Fig. 7), respectively.

**Figure 7.**
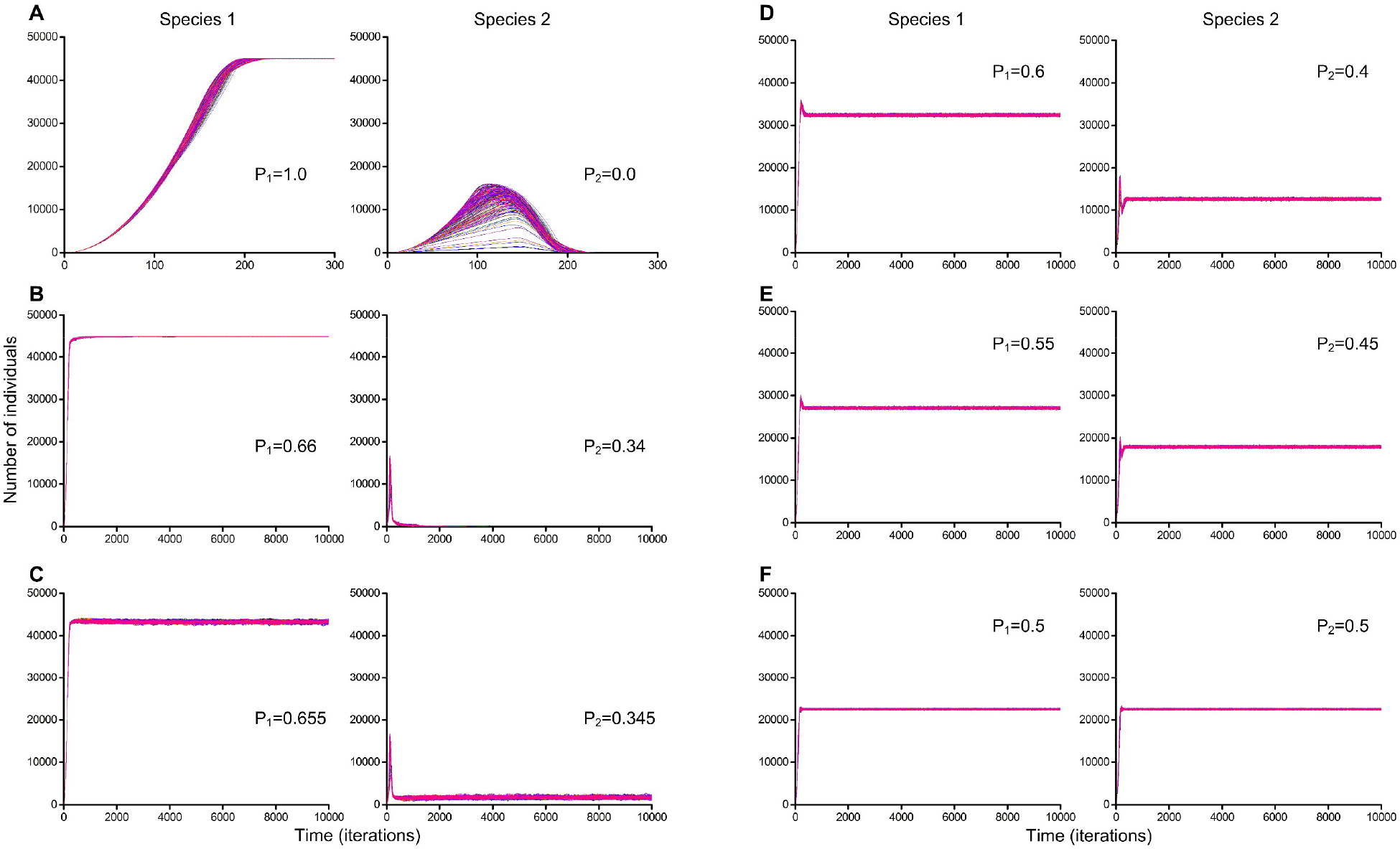
Monte Carlo simulations of two species competition with colonization of free habitat consisted of 300×300 sites. Colonization of free habitat started from single individuals of each species. The number of Monte Carlo simulations equals 200. Parameter P_1_ reflects competitiveness of the Species 1 and parameter P_2_ reflects competitiveness of the Species 2. Raw data is available on Figshare in Table S3 (https://doi.org/10.6084/m9.figshare.4903016.v1). (A, B) Competitive exclusion of the recessive Species 2 was observed in all cases. Species differences in competitiveness equal 100% (A) and 32% (B). (C-F) Stable coexistence of competing species was observed in all cases. Species differences in competitiveness equal 31% (C), 20% (D), 10% (E) and 0% (F).

#### 3.3.2 Colonization of free habitat consisted of 50×50 sites started from single individuals of each species

Here, we show experiments for the lattice consisting of 50×50 sites (Fig. 6).

#### 3.3.3 Colonization of free habitat consisted of 300×300 sites started from single individuals of each species

Further, we conducted similar experiments for the larger lattice consisting of 300×300 sites (Fig. 7).

#### 3.3.4 Colonization of free habitat consisted of 50×50 sites when the habitat is probabilistically populated by individuals on initial iteration

Next, we compared the cases in Fig. 6C with 2 individuals at the initial iteration and the cases in Fig. 8D with 625 individuals of each competing species. The habitat size had the same size – 50×50.

**Figure 8.**
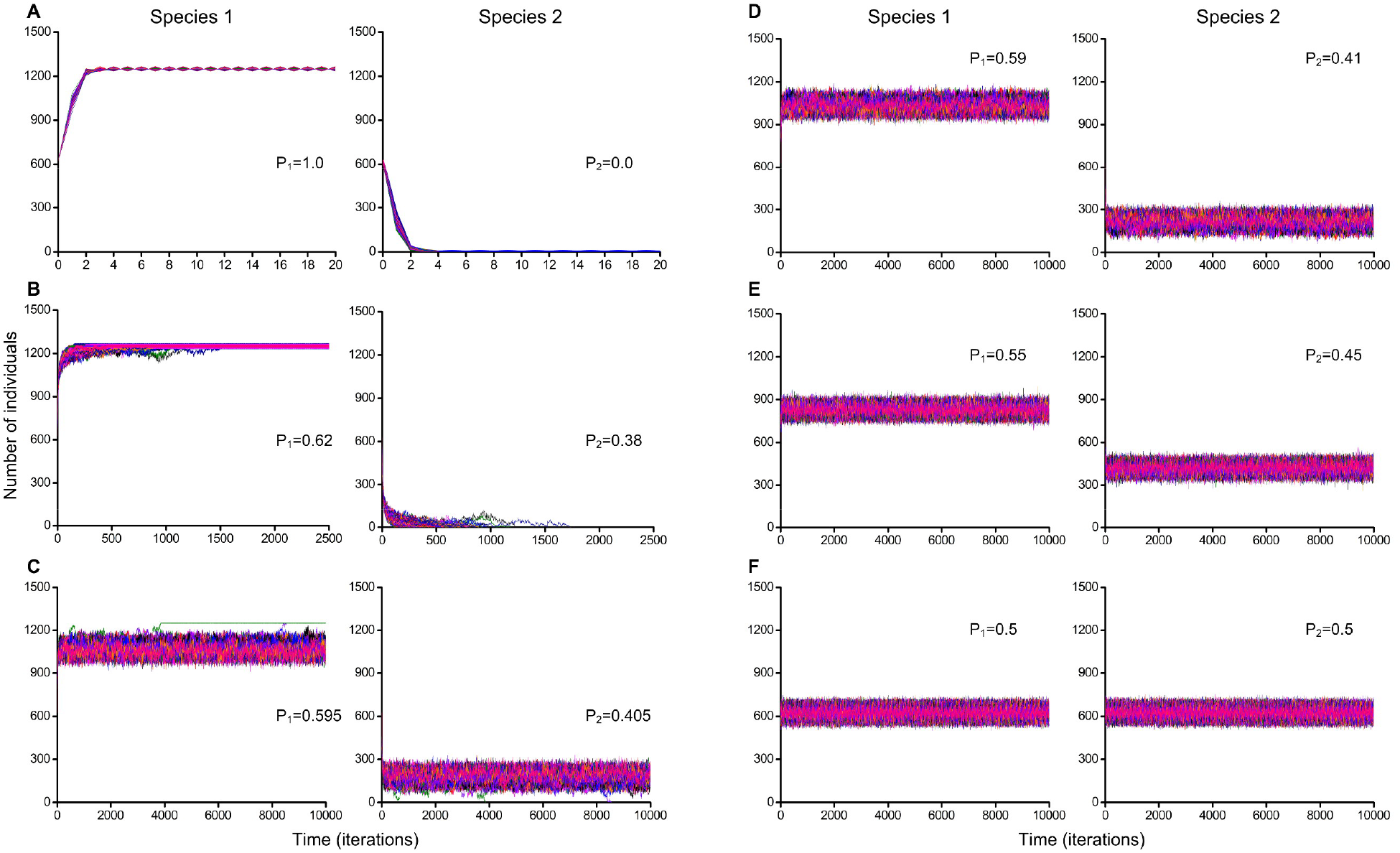
Monte Carlo simulations of two species competition when the habitat is probabilistically populated by individuals on initial iteration. Cellular automata field was probabilistically populated by 625 individuals of each of two competing species on initial iteration. The habitat consists of 50×50 microhabitats where 25% of the territory is occupied by individuals of the Species 1, 25% of the territory is occupied by individuals of the Species 2 and 50% of the territory is free. The number of Monte Carlo simulations equals 200. Parameter P_1_ reflects competitiveness of the Species 1 and parameter P_2_ reflects competitiveness of the Species 2. Raw data is available on Figshare in Table S4 (https://doi.org/10.6084/m9.figshare.4903028.v1). (A) There were 198 cases of competitive exclusion and 2 cases of competitive coexistence. Species difference in competitiveness equals 100%. (B) Competitive exclusion was observed in all cases. Species difference in competitiveness equals 24%. (C) There were 2 cases of competitive exclusion and 198 cases of competitive coexistence. Species difference in competitiveness equals 19%. (D-F) Stable coexistence of competing species was observed in all cases. Species differences in competitiveness equal 18% (D), 10% (E) and 0% (F).

#### 3.3.5 Colonization of free habitat consisted of 300×300 sites when the habitat is probabilistically populated by individuals on initial iteration

In Fig. 9 every Monte Carlo simulation consists of 200 repeated experiments with probabilistic positioning of 22500 individuals of the Species 1 and 22500 individuals of the Species 2 on the 300×300 lattice.

**Figure 9.**
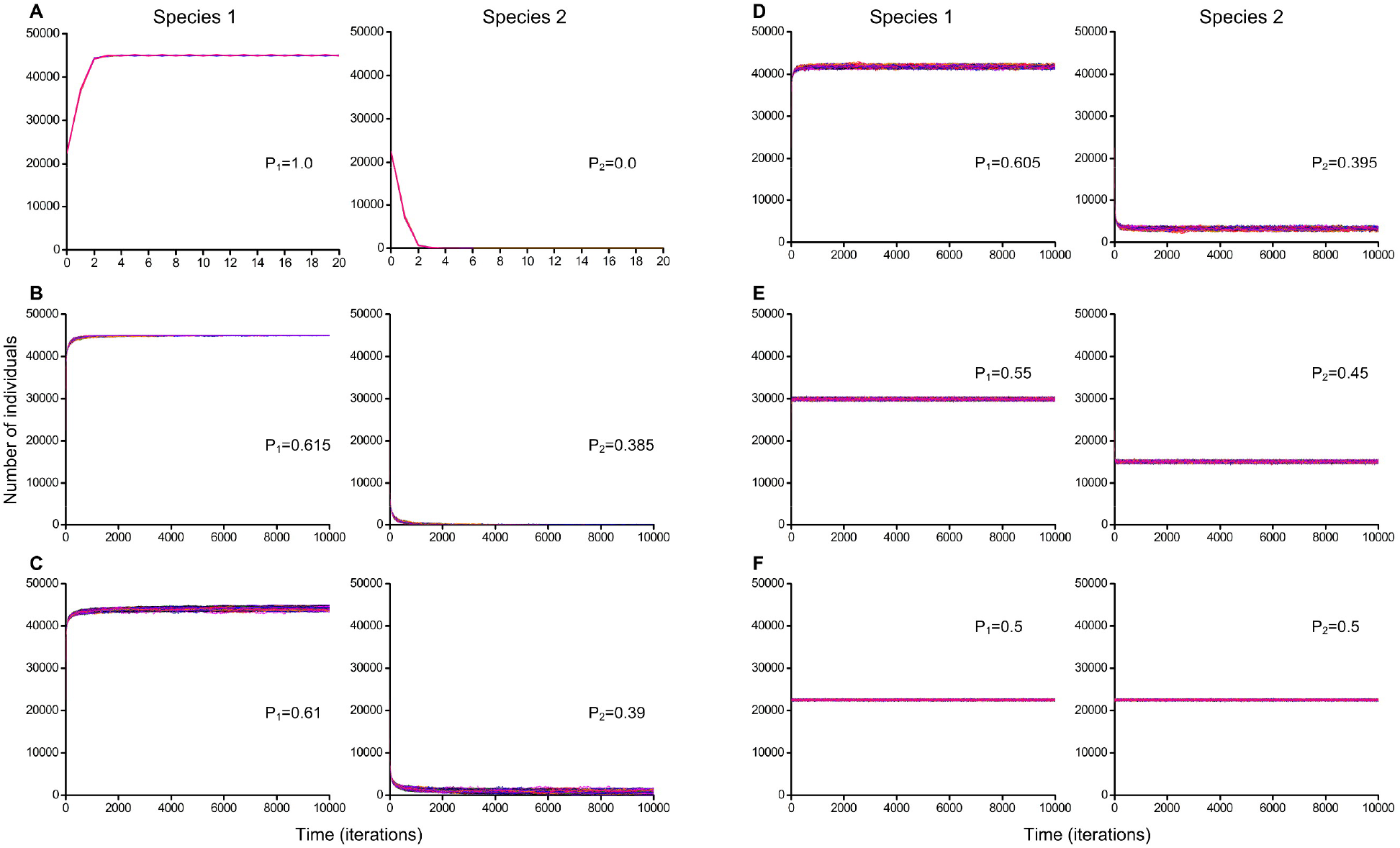
Monte Carlo simulations of two species competition when the habitat is probabilistically populated by individuals on initial iteration. Cellular automata field was probabilistically populated by 22500 individuals of each of two competing species on initial iteration. The habitat consists of 300×300 microhabitats where 25% of the territory is occupied by individuals of the Species 1, 25% of the territory is occupied by individuals of the Species 2 and 50% of the territory is free. The number of Monte Carlo simulations equals 200. Parameter P_1_ reflects competitiveness of the Species 1 and parameter P_2_ reflects competitiveness of the Species 2. Raw data is available on Figshare in Table S5 (https://doi.org/10.6084/m9.figshare.4903031.v1). (A) There were 197 cases of competitive exclusion and 3 cases of competitive coexistence. Species difference in competitiveness equals 100%. (B) There were 195 cases of competitive exclusion and 5 cases of competitive coexistence. Species difference in competitiveness equals 23%. (C-F) Stable coexistence of competing species was observed in all cases. Species differences in competitiveness equal 22% (C), 21% (D), 10% (E) and 0% (F).

In the case shown in Fig. 9A there are 3 cases of competitive coexistence as a result of implementation of the mechanism of avoiding a direct conflict of interest, which was demonstrated in Fig. 8A and Supplemental Video S4.

#### 3.3.6 The outcome of competition between two completely competing species when relative competitiveness depends on a number of individuals locally competing for a free microhabitat

Fig. 10 shows that 10% of differences in competitiveness did not prevent competitive coexistence in the conditions of the studied cooperative effects of competitiveness dependence on the number of locally competing individuals. With 20% difference there was a competitive exclusion of the recessive species. With the equal competitiveness the competing species stably coexist.

**Figure 10.**
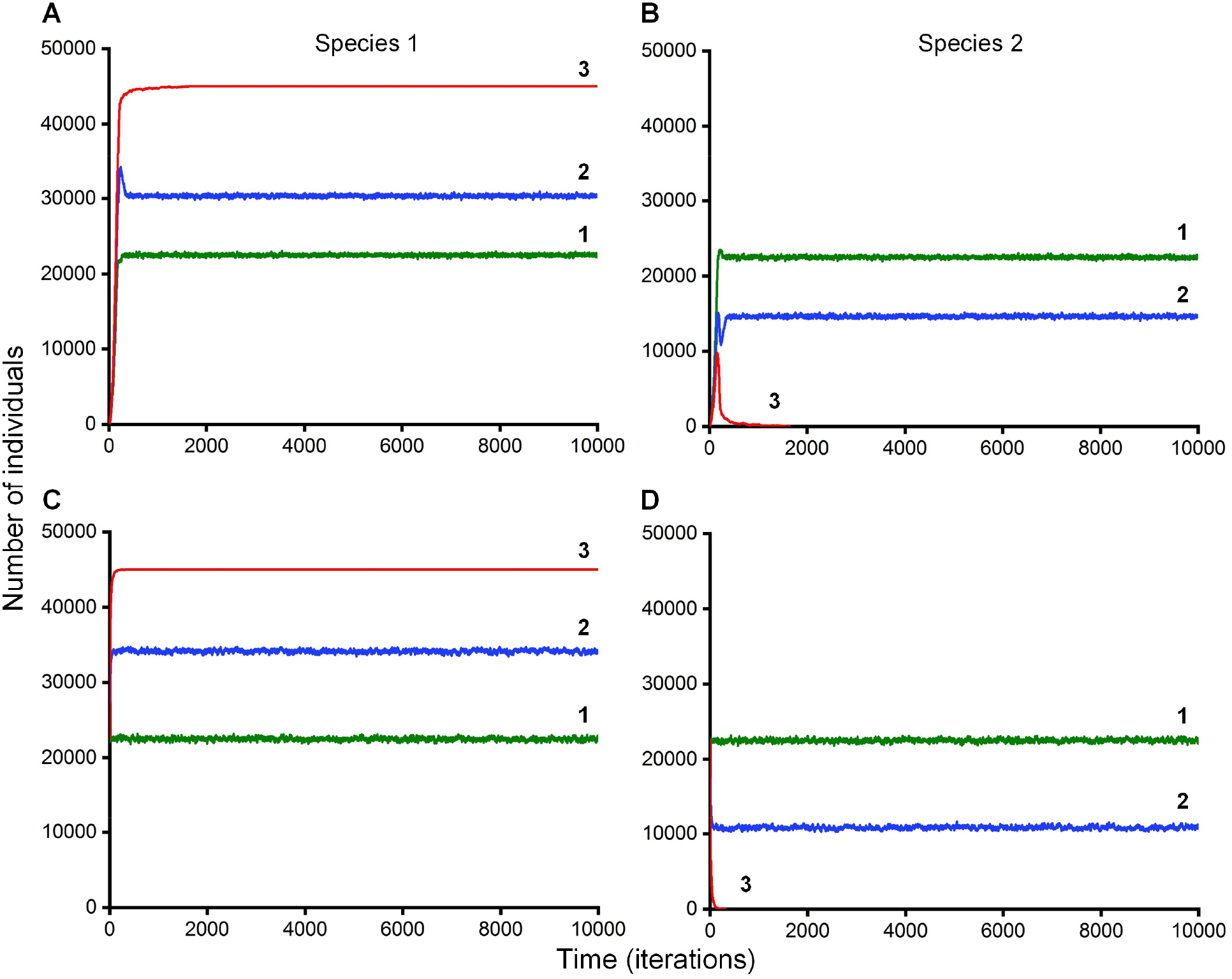
The outcome of competition between two completely competing Species 1 (sp.1) and Species 2 (sp. 2) when relative competitiveness P depends on a number of individuals locally competing for a free microhabitat. Initial competitiveness of the species: “1” (green curves) P_sp.1_ = 0.5, P_sp.2_ = 0.5; “2” (blue curves) P_sp.1_ = 0.55, P_sp.2_ = 0.45; “3” (red curves) P_sp.1_ = 0.6, P_sp.2_ = 0.4. The lattice size is 300×300 in all cases. **A** and **B –** the initial pattern of the habitat is populated by two single individuals of two competing species; the **numerical superiority per one individual** in a local conflict of interests increases the competitiveness of the respective species in a given conflict by **8%**. **C** and **D –** the initial pattern of the habitat is randomly populated by individuals – 25% by the Species 1, 25% by the Species 2 and 50% of the habitat remains free; the **numerical superiority per one individual** in a local conflict of interests increases the competitiveness of the respective species in a given conflict by **6%**.

## 4 DISCUSSION

### 4.1 On preliminary verification of the main hypothesis (see 3.1 in Results)

The case of competitive exclusion (Fig. 4A and Supplemental Video S1) closely reproduces dynamics from the Gauze’s experiments in which *Paramecium aurelia* drives out *Paramecium caudatum*, when they were cultivated in the mixed population with minimal nutrition (Gause 1934). In our case the complete competitors cannot coexist and only the fittest Species 1 survives. Collision of population waves of the two species in the habitat leads to annihilation of population waves of the Species 2. In Fig. 4B and Supplemental Video S2 we show how two completely competing species coexist despite 29% difference in competitiveness. In the case when complete competitors have the same competitiveness, they coexist with close numbers of individuals (Fig. 4C and Supplemental Video S3). This dynamic shows diffusion-like behavior of colliding population waves. It demonstrates the special case of ecological equivalence of competing species which we consider as ecological neutrality. In contrast to the unified neutral theory of biodiversity and biogeography (Hubbell 2001), our models explicitly take into account local interactions between individuals.

Figure 4B and Supplemental Video S2 demonstrate a possibility of coexistence of competing species with 29% difference in competitiveness. A procedure of determination of a winner in each conflict of interest between individuals of competing species has a probabilistic nature (Fig. 1). This result confirms the main hypothesis of the article.

### 4.2 On contribution of stochastic fluctuations in coexistence in small ecosystems (see 3.2 in Results)

On the 5×5 lattice competitive exclusion of one of the species was observed already at 11-th iteration (Fig. 5A). On the 6×6 lattice species coexisted 504 iterations (Fig. 5B). On the 7×7 lattice the competitive exclusion required 586 iterations (Fig. 5C). On the 8×8 lattice the species coexisted during 5000 iterations (Fig. 5D). The obtained results demonstrate that the smaller the size of the lattice we have, the more stochastic fluctuations of competition outcomes appear in favor of one of the species. This individual-based model reproduces a phenomenon of the neutral genetic drift (Turner and Miller 2015), based on stochastic fluctuations similar to the Moran process (Moran 1958). In small ecosystems, even at equal competitiveness of both species, stochastic fluctuations may drive one of the species to extinction. A simple analogy with symmetrical coin – if it is tossed only several times, it can show an occasional repetition of the same side. However, if this procedure will be repeated one million times, then the result will differ slightly from 50% for the both sides of the coin. Increasing of the lattice size leads to increasing in the number of cases of local competition, and, as a consequence, the contribution of probabilistic fluctuations in favor of one of the species is reduced, and the revealed phenomena become more natural and lawful.

### 4.3 On colonization of free habitat consisted of 50×50 sites started from single individuals of each species (Monte Carlo simulations) (see 3.3.1, 3.3.2 in Results)

In the case, when competing species had a maximum difference in competitiveness (P_1_ = 1.0; P_2_ = 0.0), then the Species 1 always outcompeted the Species 2 (Fig. 6A). This case does not contradict with the classical formulations of the competitive exclusion principle. In another case, when P_1_ = 0.65 and P_2_ = 0.35 there were 78 cases of competitive exclusion and 122 cases of competitive coexistence (Fig. 6B). When P_1_ = 0.645 and P_2_ = 0.355, i.e. the difference in competitiveness is 29%, competing species stably coexisted in all cases despite of random initial positioning of individuals of competing species in the habitat (Fig. 6C). At smaller differences in competitiveness the competing species also coexisted and their difference in number of individuals was smaller (Fig. 6C-F). These cases (Fig. 6C-F) demonstrate the validity of our hypothesis that two completely competing species can coexist with smaller difference in competitiveness despite of random initial positioning of individuals of competing species in the habitat.

### 4.4 On colonization of free habitat consisted of 300×300 sites started from single individuals of each species (see 3.3.3 in Results)

Further, we conducted similar experiments for the larger lattice consisting of 300×300 sites (Fig. 7). We showed that due to increasing of the lattice size, competing species could coexist regardless initial placement of individuals with larger competitiveness difference of 31% (Fig. 7C). On the lattice of 50×50 sites it is 29%. Thus, the influence of stochastic fluctuations in result of competitive interactions has been decreased noticeably. In the case demonstrated on Fig. 7F, the species have identical characteristics and have little difference in number after occupation of the habitat.

In Fig. 8 every Monte Carlo simulation consists of 200 repeated experiments with probabilistic positioning of 625 individuals of each species on the 50×50 lattice. An example of the initial pattern is shown in Fig. 3B. We discovered realization of a case of coexistence of complete competitors in 2 cases from 200 Monte Carlo experiments (Fig. 8A, Supplemental Video S4, Raw data in Table S4 - https://doi.org/10.6084/m9.figshare.4903028.v1). As a result of random initial location of individuals on the lattice, a particular pattern was realized that allowed the recessive species to avoid a direct competition for resources with individuals of the dominant species. However, in such rare situation, the number of individuals of the recessive Species 2 was very small and fluctuated periodically between 1 and 6 – 1, 6, 1, 6, etc. This recessive species was on the verge of extinction, although it did not disappear – one individual of recessive species successfully opposes to 1246 individuals of the dominant species.

Increasing the habitat size from 50×50 to 300×300 resulted in an increase in the number of coexistence cases, – species always coexisted with differences in competitiveness equal to 29% (Fig. 6C) and 31% (Fig. 7C), respectively.

### 4.5 On colonization of free habitat consisted of 50×50 sites when the habitat is probabilistically populated by individuals on initial iteration (see 3.3.4 in Results)

In Fig. 6C the maximum value of the parameter P at which the species stably coexist equals 0.29 (29% difference in competitiveness) and in Fig. 8D the parameter P equals 0.18 (18% difference in competitiveness). Thus, when on initial iteration the habitat was populated only by 2 individuals there were the larger number of cases of competitive coexistence than when 50% of the habitat was initially populated (Figs. 3, 6, 8). At the identical sizes of the habitat, the species coexisted better when their competition began with two individuals at the initial iteration.

### 4.6 On colonization of free habitat consisted of 300×300 sites when the habitat is probabilistically populated by individuals on initial iteration (see 3.3.5 in Results)

Increasing of the habitat size has led to increasing the number of cases of coexistence of competing species in conditions of direct competition for resources (Figs. 8D, 9C). In Fig. 8D species coexisted in all 200 cases at P = 0.18 (18% difference in competitiveness), and in Fig. 9C at P = 0.22 (22% difference in competitiveness).

Figs. 8C and 9B demonstrate cases of competitive exclusion when species had minimal differences in competitiveness. Decreasing the difference in competitiveness at 1% led to stable coexistence in all cases (Figs. 8D and 9C).

### 4.7 The outcome of competition between two completely competing species when relative competitiveness depends on a number of individuals locally competing for a free microhabitat (see 3.3.6 in Results)

The results are shown in Fig. 10. In case (A), ecologically identical species coexisted with close numbers, in case (B) the species coexisted contrary to the known formulations of the competitive exclusion principle; in case (C), the recessive species was replaced by the dominant one. The cooperative dependence on the number of locally competing individuals left the main conclusions unchanged. The main hypothesis of the article has been confirmed - two completely competing species may coexist with smaller than 100% difference in competitiveness regardless of random initial location of individuals of competing species in the habitat. In the experiments with cooperative dependence complete competitors stably co-existed at 10% difference in basic competitiveness. With 20% difference, there was a competitive exclusion.

### 4.8 General discussion

Here we have discovered a new fact that two aggressively spreading competitors that are identical consumers can coexist stably on the same limiting resource in the same limited, stable and uniform habitat when one species has some competitive advantage over others, and all other characteristics are equal.

We have found that when colonization of free habitat started from a single individual of each species, then the complete competitors coexisted up to 31% of their difference in competitiveness and when on initial stage half of the territory was probabilistically occupied, the complete competitors coexisted up to 22% of their difference in competitiveness. With additional cooperative effects the competitive coexistence also took place, albeit with fewer differences in basic competitiveness (10%). These results may be considered as the answers on the question on the “limiting similarity” - how much species must differ in their functional traits to implement competitive exclusion (Hutchinson 1959, Macarthur and Levins 1967). We managed to anatomize competitiveness by separating food specialization from priority effects and functional traits. We consider that priority effects and some particular species traits may be mechanisms of the difference in competitiveness modelled here. A priority effect may be the impact that a particular species can have on community development due to prior arrival at a site, for example, due to inhibitory priority effects when a species that arrives first at a site negatively impacts a species that arrives later by reducing the availability of space or resources (Fukami et al. 2005). A distinguishing feature of the ability to capture a resource, may also be a consequence of a specific functional eco-physiological traits - such as rate and strength of growth (Fukami et al. 2005). The revealed coexistence of complete competitors with competitiveness inequality is carried out regardless of random initial location of individuals in the ecosystem, and, consequently, the main hypothesis of this study has been confirmed.

The conditions of our experiments were extremely unfavourable for competitive coexistence. Earlier we found *two* deterministic individual-based mechanisms of competitive coexistence (Kalmykov and Kalmykov 2013, 2015b). *The first coexistence mechanism* is based on free resource gaps which help to eliminate direct conflicts of interest between competing species, and as result colliding population waves of different competing species interpenetrate through each other like soliton waves in physical systems (Kalmykov and Kalmykov 2013). A possible mechanism of appearance of such gaps is moderate reproduction which was modelled through a hexagonal rosette-like cellular automata neighbourhood. *The second coexistence mechanism* is based on timely recovery of the limiting resource, its spatio-temporal allocation between competitors and limitations of habitat size (Kalmykov and Kalmykov 2015b). This mechanism allows complete competitors coexist in spite of using standard hexagonal cellular automata neighbourhood which models aggressive propagation without gaps in population waves. However, this mechanism of indefinite coexistence was limited by the habitat size and initial location of individuals on the lattice. In any case, the principle must always be right and if there are exceptions, then either the principle is not true, or its wording is not correct.

A large number of studies have been devoted to finding mechanisms that prevent the implementation of the competitive exclusion principle. There were found more than 120 (Palmer 1994) of such mechanisms (Hutchinson 1961, Hastings 1980, Wellborn 2002, Dollhopf et al. 2003, Nowak 2006, Bennett and Bever 2009). However, identification of factors hindering the implementation of the principle of competitive exclusion had little effect on the formulation of the principle itself. At the same time, formulations of the principle gradually became more and more stringent:
- Survival of the fittest (Spencer 1864);
- Complete competitors cannot coexist (Hardin 1960);
- *n* species require at least *n* resources to ensure indefinite and stable equilibrium coexistence in a homogeneous environment (MacArthur and Levins 1964, Narwani et al. 2009);
- No stable equilibrium can be attained in an ecological community in which some r of the components are limited by less than r limiting factors (Levin 1970);
- Two populations (species) cannot long coexist if they compete for a vital resource limitation of which is the direct and only factor limiting both populations (Darlington 1972).
- Given a suite of species, interspecific competition will result in the exclusion of all but one species. Conditions of the Principle: (1) Time has been sufficient to allow exclusion; (2) The environment is temporally constant; (3) The environment has no spatial variation; (4) Growth is limited by one resource; (5) Rarer species are not disproportionately favoured in terms of survivorship, reproduction, or growth; (6) Species have the opportunity to compete; (7) There is no immigration (Palmer 1994).

To carry out a rigorous verification of the principle, we made an attempt to find a mechanism for competitive coexistence under the extremely strict conditions of interspecific competition. In our experiments we excluded a possibility of implementing numerous mechanisms that prevent implementation of the principle (Palmer 1994).

The revealed mechanisms of competitive coexistence violate the listed formulations of the competitive exclusion principle and as result we have reformulated it as follows (Kalmykov and Kalmykov 2013, 2015b, Kalmykov and Kalmykov 2016):

#### (Definition 1)

*If each and every individual of a less fit species in any attempt to use any limiting resource always has a direct conflict of interest with an individual of a most fittest species and always loses, then, all other things being equal for all individuals of the competing species, these species cannot coexist indefinitely and the less fit species will be excluded from the habitat in the long run*.

These strict clarifications in the formulation of the principle demonstrate a low probability of the realization of this principle in nature because the need to comply with all formulated conditions makes the competitive exclusion rather a rare event. The obvious rarity of implementation the competitive exclusion principle removes the paradox of biodiversity because it eliminates the contradiction between the principle and the observed natural facts. In addition, we generalized the reformulated principle Def.1, setting out conditions under which one competitor is able to displace all others (Kalmykov and Kalmykov 2015b, Kalmykov and Kalmykov 2016):

#### (Definition 2)

*If a competitor completely prevents any use of at least one necessary resource by all its competitors, and itself always has access to all necessary resources and the ability to use them, then, all other things being equal, all its competitors will be excluded*

This formulation allows us to interpret different mechanisms of coexistence in terms of availability of access to necessary resources. It helps us to understand a threat to biodiversity that may arise if one competitor will control use of a necessary resource of an ecosystem. Nowadays, humankind is becoming such a global competitor for all living things (Vitousek et al. 1997, Cincotta et al. 2000, Ehrlich and Ehrlich 2008). Overexploitation of limited necessary resources in result of unbridled human population growth causes a problem which is known as the ‘tragedy of the commons’ (Hardin 1959, 1968).

Starting from the revealed mechanisms of competitive coexistence and competitive exclusion and reasoning by contradiction regarding the generalized competitive exclusion principle (Def. 2), we formulated *a principle of competitive coexistence*, which is valid for any number of potential resource competitors of any nature (Kalmykov and Kalmykov 2015b, Kalmykov and Kalmykov 2016):

#### (Definition 3)

*If competitors always have access to all necessary resources, and all have the ability to use them, then, all other things being equal and without a global catastrophe, they will coexist indefinitely.*

Under a global catastrophe we mean a disaster, a fatal disease pandemic, a devastating invasion of predators or herbivores, a mutual destruction. Indefinite competitive coexistence implies timely access of all competitors to all necessary resources. Such access is the necessary condition for biodiversity conservation and sustainable development.

## 5 CONCLUSIONS

Here, along with the coexistence of complete competitors differing in competitiveness, we demonstrated the cases of the classic competitive exclusion. On the other hand, if complete competitors have equal competitiveness, then on lattices with not less than 8×8 sites, they stably coexist with close numbers of individuals, demonstrating “neutrality”. Consequently, our approach unifies competitive exclusion (“niche”), neutrality and coexistence of complete competitors in one model.

We hope that our axiomatic ecosystem theory contributes to raising the level of theoretical ecology to the level of theoretical physics. The ‘bottom-up’ individual-based approach to hypotheses verification we implemented as an automatic deductive inference and it is a promising method of transparent artificial intelligence. Our method of automatic deductive inference opens up prospects for a deeper study of mechanisms of complex systems and for developments in the field of transparent artificial intelligence. Conventional logic is based only on linear sequential operations. Automatic deductive inference based on cellular automata is implemented hyperlogically, i.e. logical operations are carried out simultaneously, in parallel and interconnected (through the cellular automata neighborhood). Behavior of all micro-objects (micro-ecosystems) integrated holistically at each iteration with each other and with the macrosystem (macro-ecosystem). This leads to the disappearance of fundamental differences between logic and image, since each individual iteration of the logical evolution of cellular automata is implemented as a two-dimensional logical pattern that stitches the logical events underlying dynamics the macro-ecosystem into a single whole. This cellular automata hyperlogics is the basis of our method of transparent artificial intelligence.

The presented research demonstrated the possibility of transparent multi-level modeling of complex systems by combining the axiomatic theory of ecosystem with the formalism of logical cellular automata. This ecosystem-inspired modeling of a complex system is based on a transparent artificial intelligence and opens up great prospects for a variety of theoretical and applied fields.

## ACKNOWLEDGEMENTS

The authors are grateful to Lev Naumov for permission to use his source code of the John Conway’s Game of Life as a starting point for designing of the programs. We thank Markus Dahlem, Jörg Weimar and Cody Dey for their detailed comments and suggestions for the earlier versions of the manuscript. This research was partially supported by the Russian Foundation for Basic Research (grant No. 16-31-00516).

## COMPETING INTERESTS

The authors declare there are no competing interests.

## AUTHORSHIP

Both authors conceived and designed the experiments, contributed analysis tools, analyzed the data, wrote the paper. L. V. K. performed the experiments, created programs and movies.

## DATA ACCESSIBILITY

Supplemental Videos S1-S4 are available online.

Raw data for Figure 5 is available on Figshare in Table S1 (https://doi.org/10.6084/m9.figshare.4903013.v1).

Raw data for Figure 6 is available on Figshare in Table S2 (https://doi.org/10.6084/m9.figshare.4902986.v1).

Raw data for Figure 7 is available on Figshare in Table S3 (https://doi.org/10.6084/m9.figshare.4903016.v1).

Raw data for Figure 8 is available on Figshare in Table S4 (https://doi.org/10.6084/m9.figshare.4903028.v1).

Raw data for Figure 9 is available on Figshare in Table S5 (https://doi.org/10.6084/m9.figshare.4903031.v1).

Raw data for Figure 10 is available on Figshare in Table S6 (https://doi.org/10.6084/m9.figshare.7144595.v1)

## Appendix S1: The source codes of the individual-based cellular automata Model 1 and Model 2 of an ecosystem with two resource competitors

These four source codes were written in C++ programming language and tested in Microsoft Visual C++ Studio 2015. The programs allow you to visually examine all cases of competition presented in the article. It is possible to change habitat sizes and parameters of the species competitiveness. The number of individuals of each competing species is recorded in a file at each iteration and can be analyzed further. The programs carry out global behavior modeling from local rules. It allows to implement transparent individual-based modeling approach based on the cellular automata logic. The transparency means not only that individuals are presented in the model, but also that we can control the dependence of each individual’s behavior on its local conditions. This allowed us to study mechanisms of interspecific competition in a quantitative and the most complete way, taking into account the relative competitiveness of each and every individual. This ‘bottom-up’ individual-based approach to hypotheses verification is implemented as an automatic deductive inference and it is a promising method of the transparent artificial intelligence.

**Source code 1.**
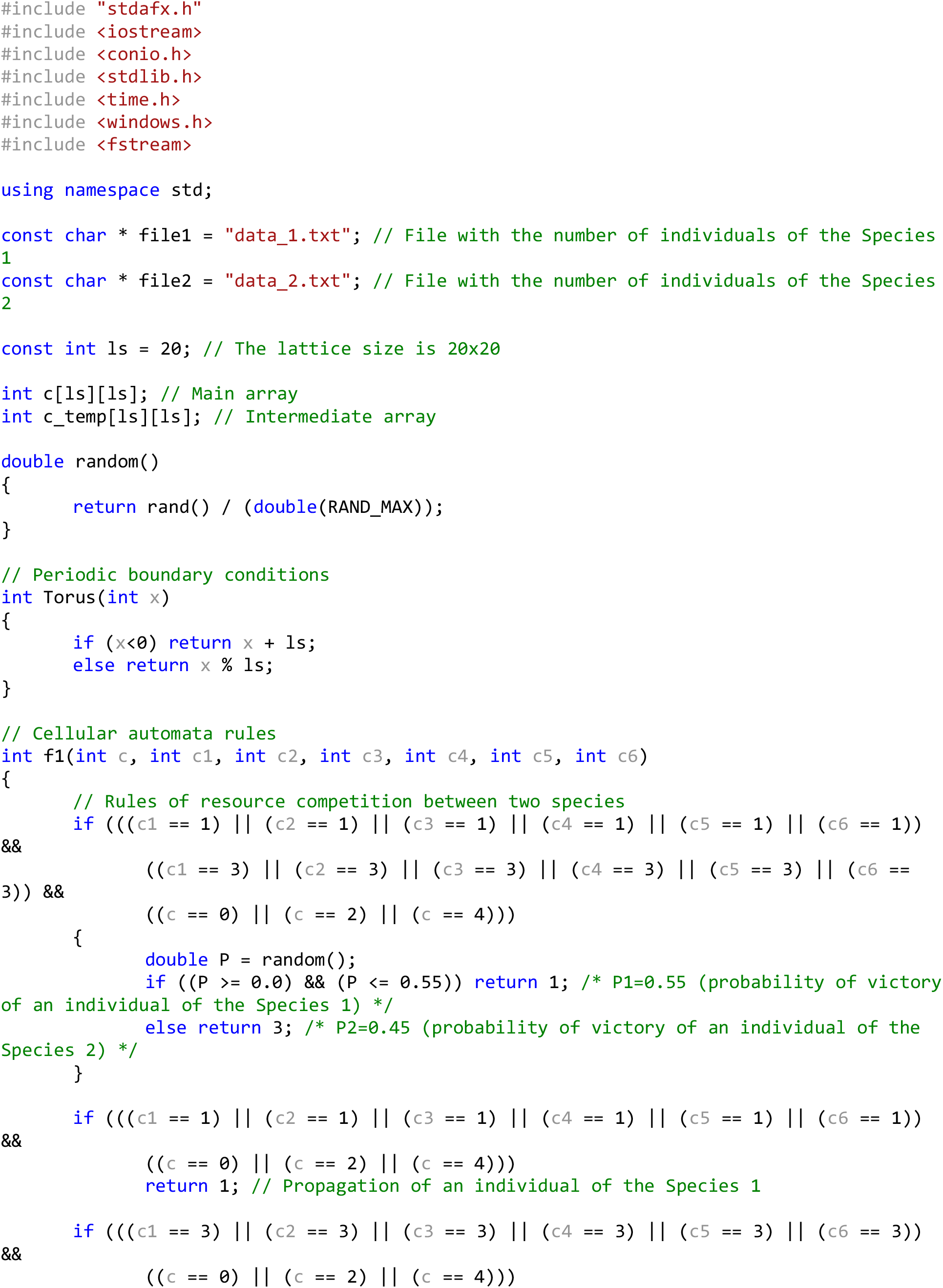
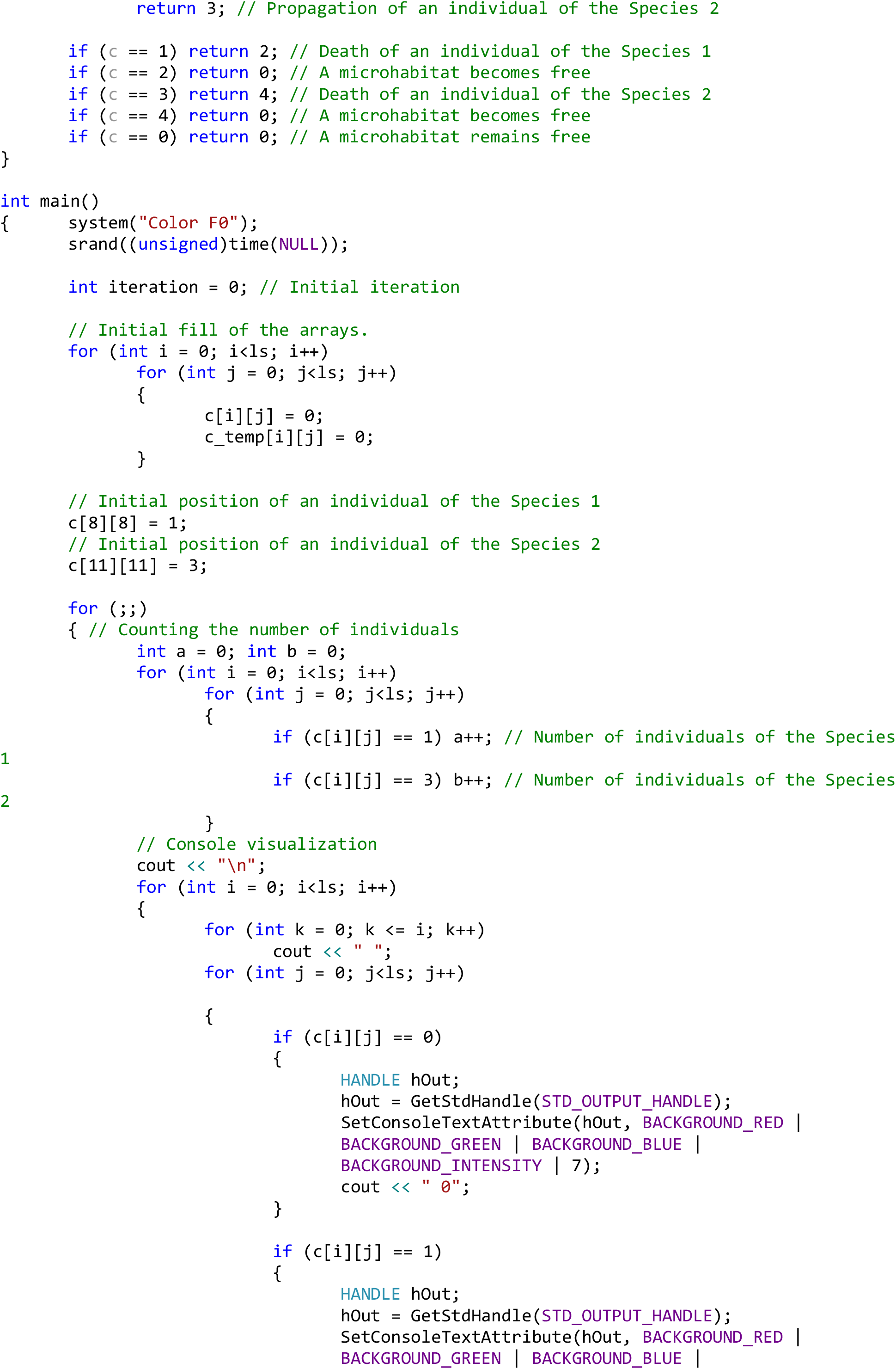
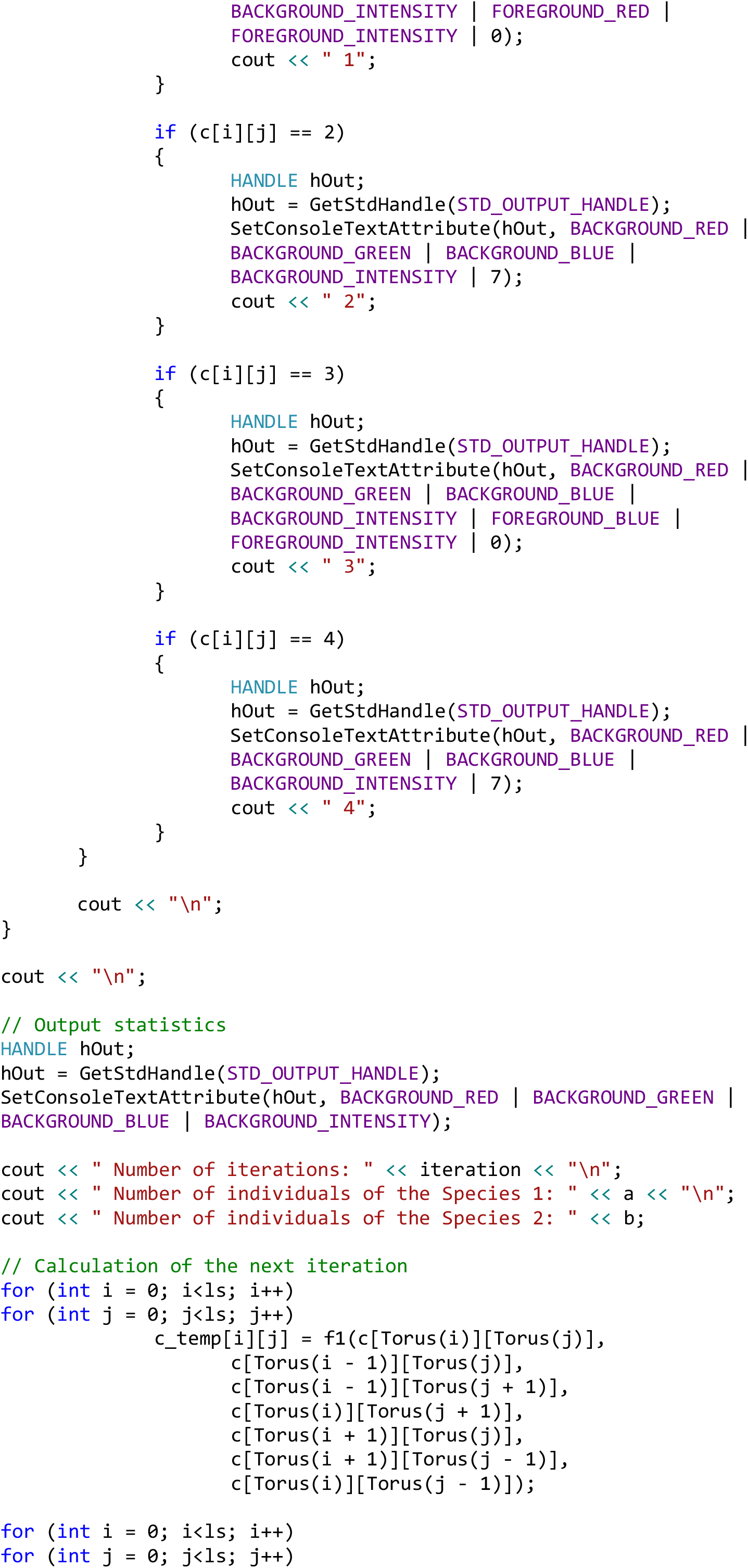
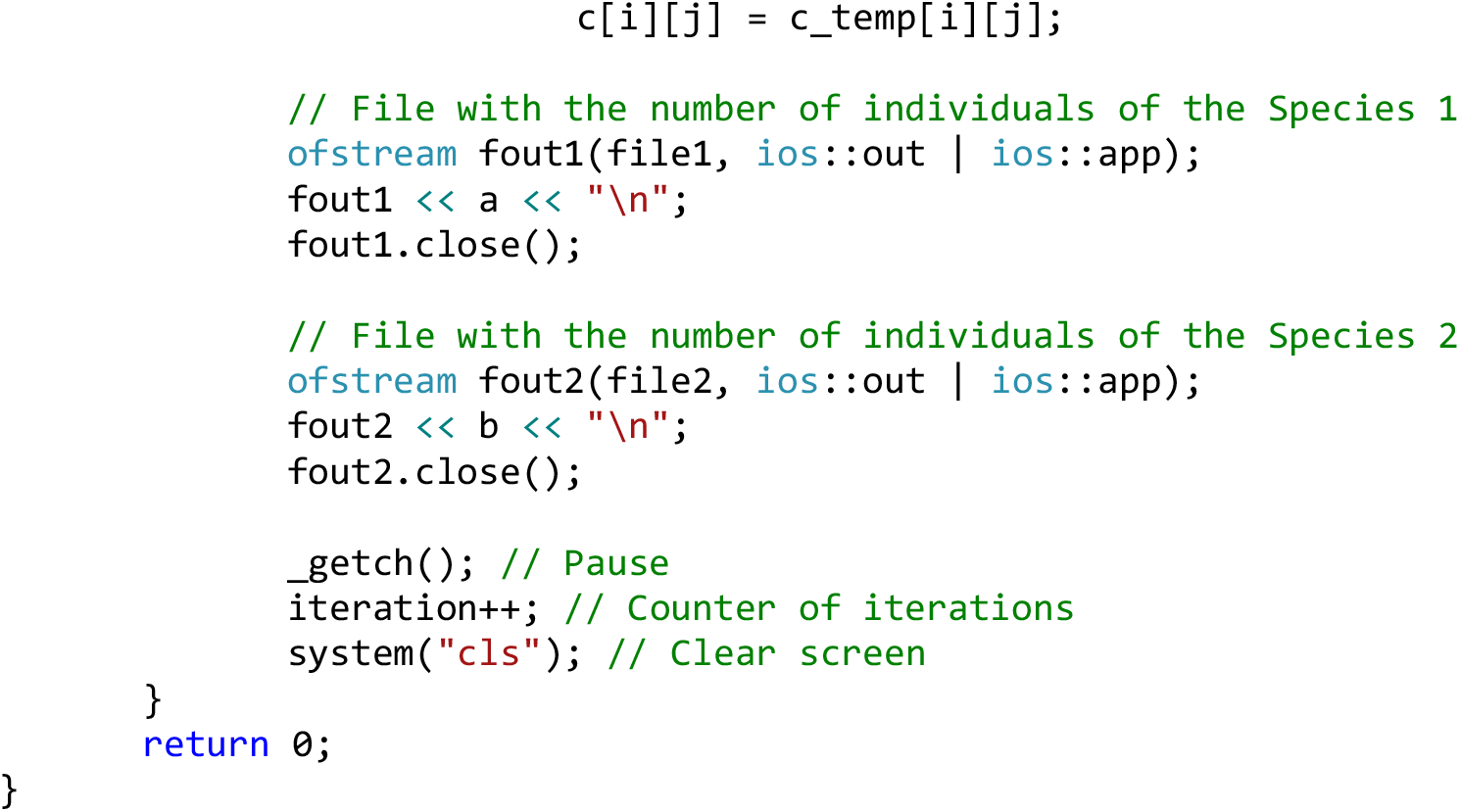
Model 1. Coexistence of complete competitors with 10% difference in fitness without cooperative dependence of fitness on the number of locally competing individuals. The initial pattern of the habitat is populated by single individuals of two competing species.

**Source code 2.**
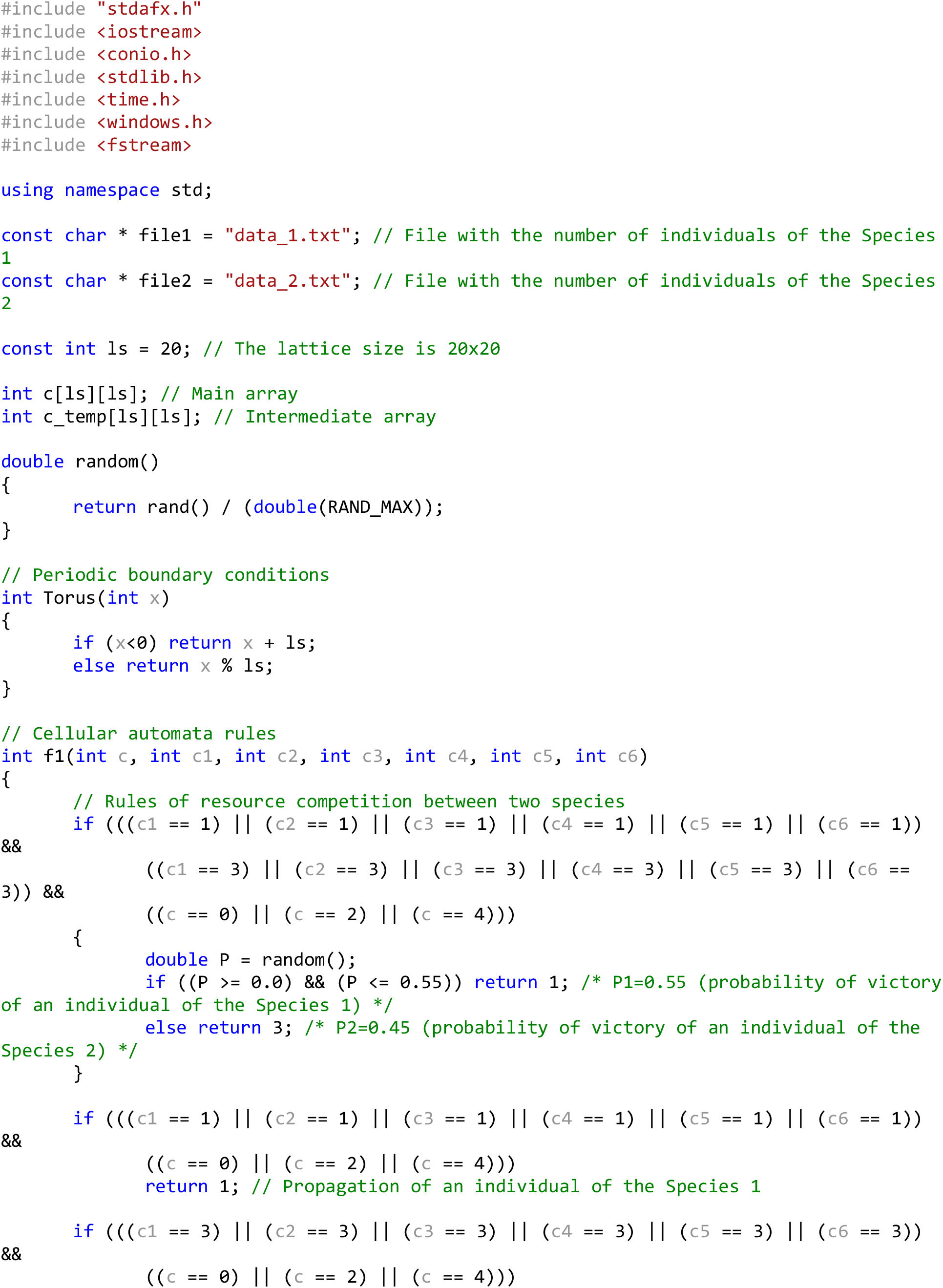
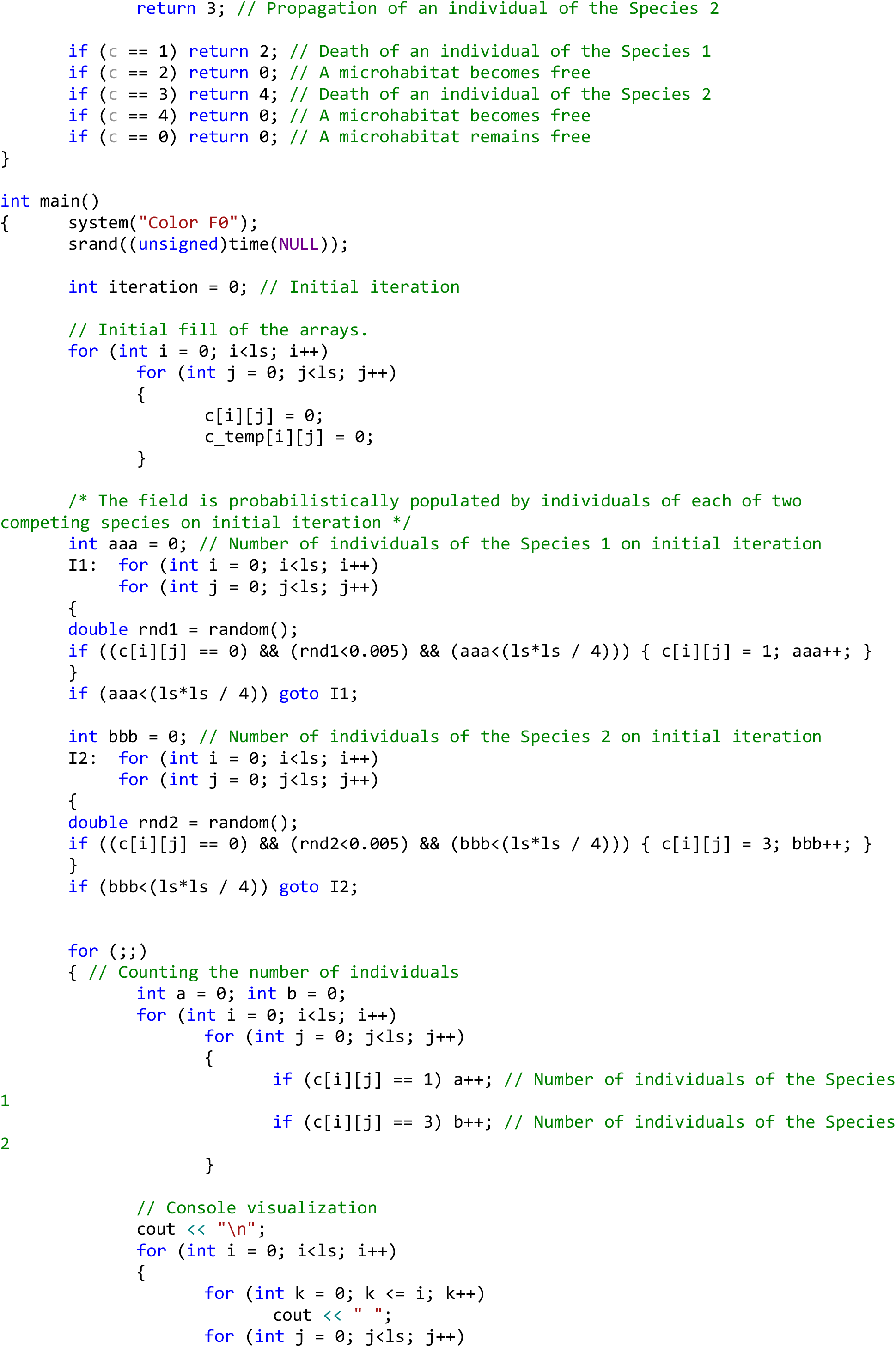
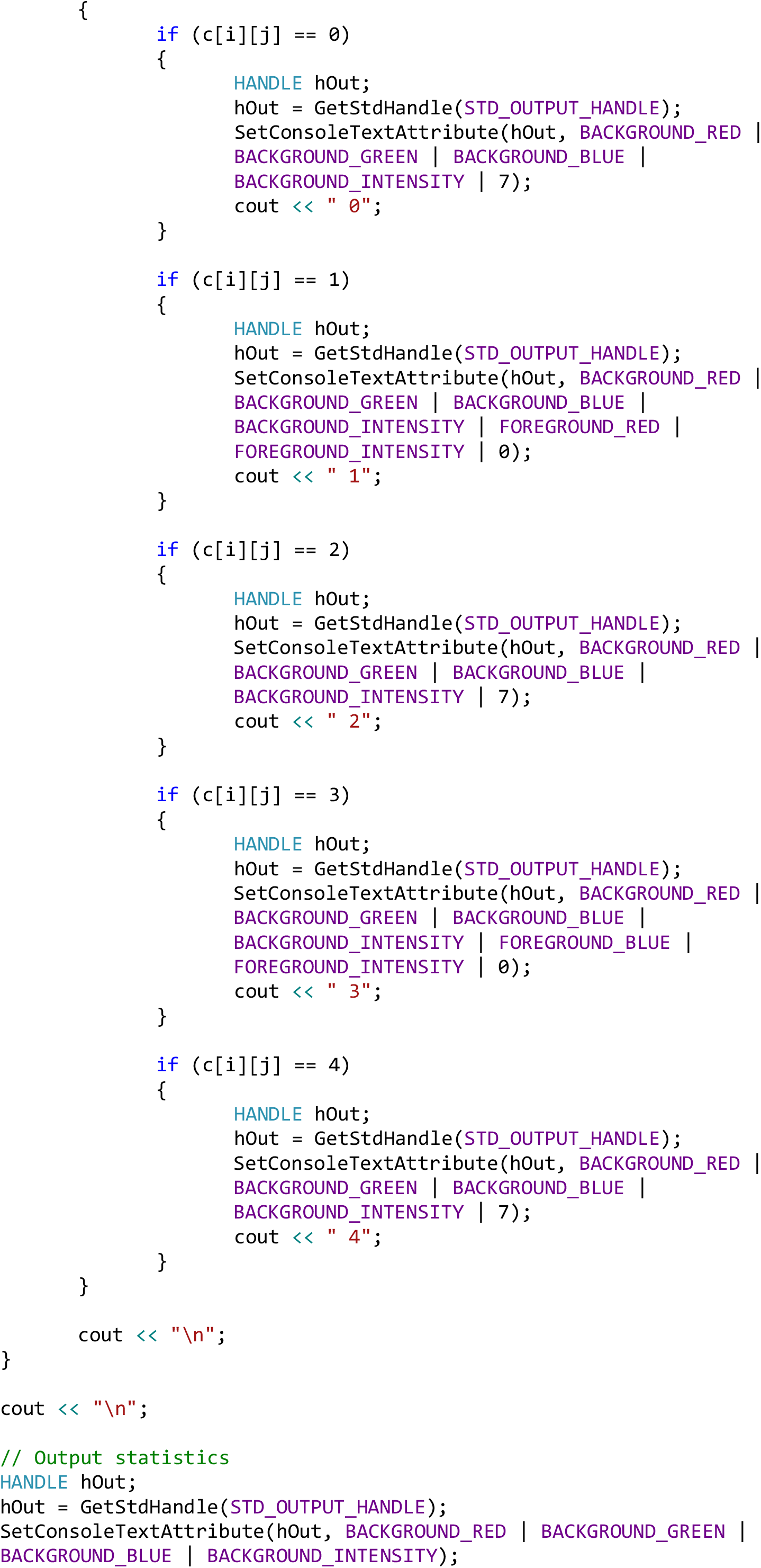
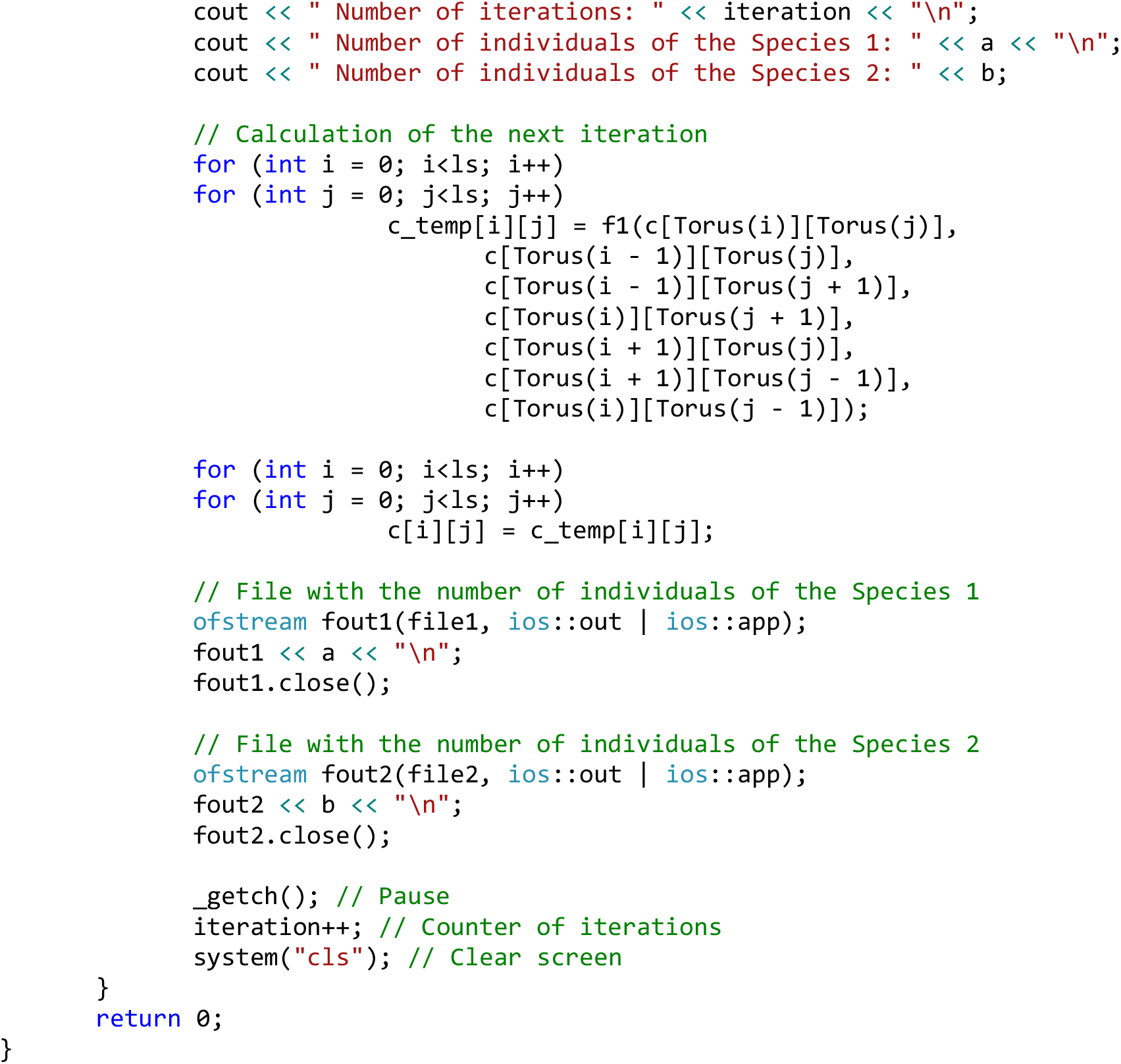
Model 1. Coexistence of complete competitors with 10% difference in fitness without cooperative dependence of fitness on the number of locally competing individuals. The initial pattern of the habitat is randomly populated by individuals - 25% by the Species 1, 25% by the Species 2 and 50% of the habitat remains free.

**Source code 3.**
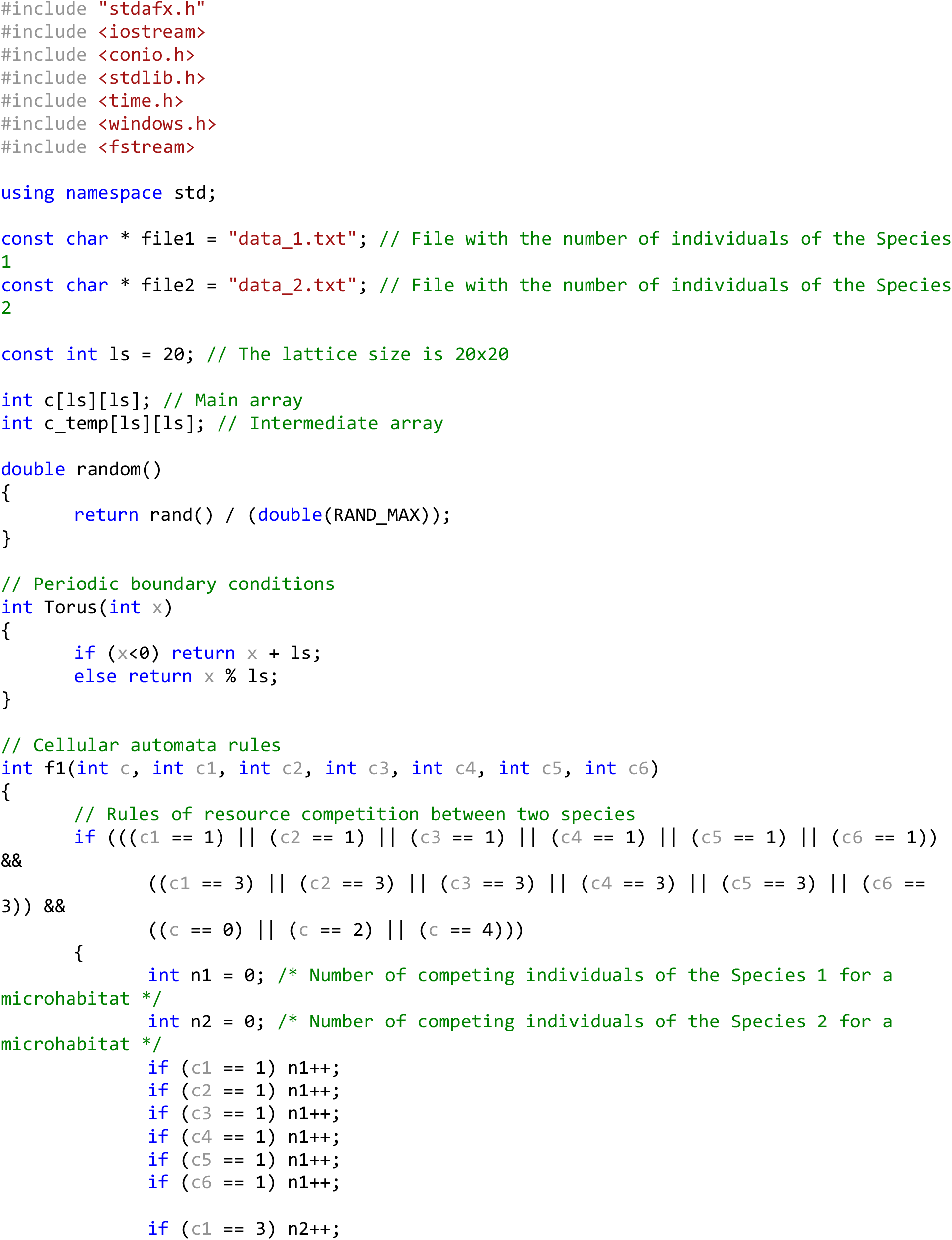
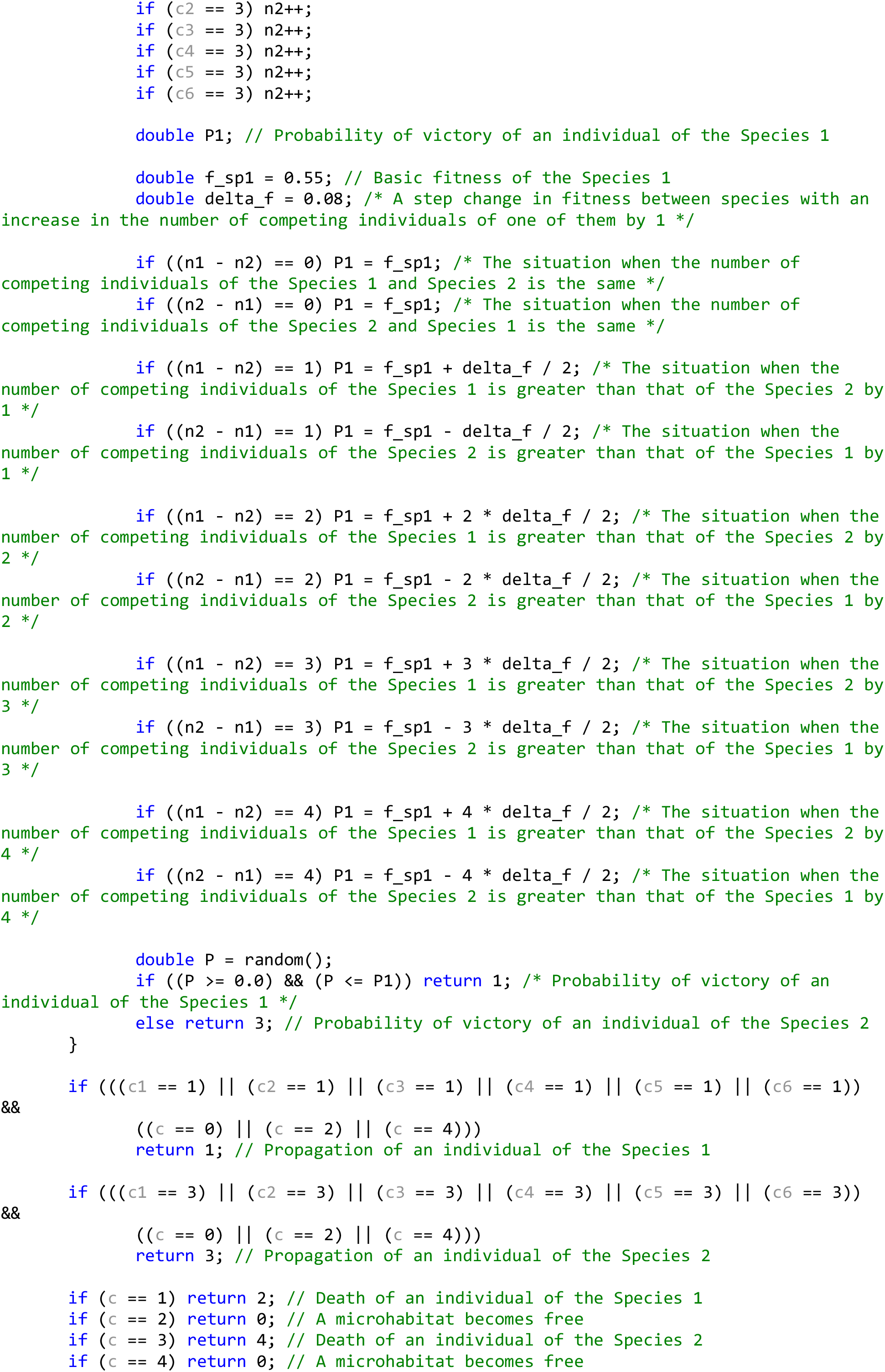
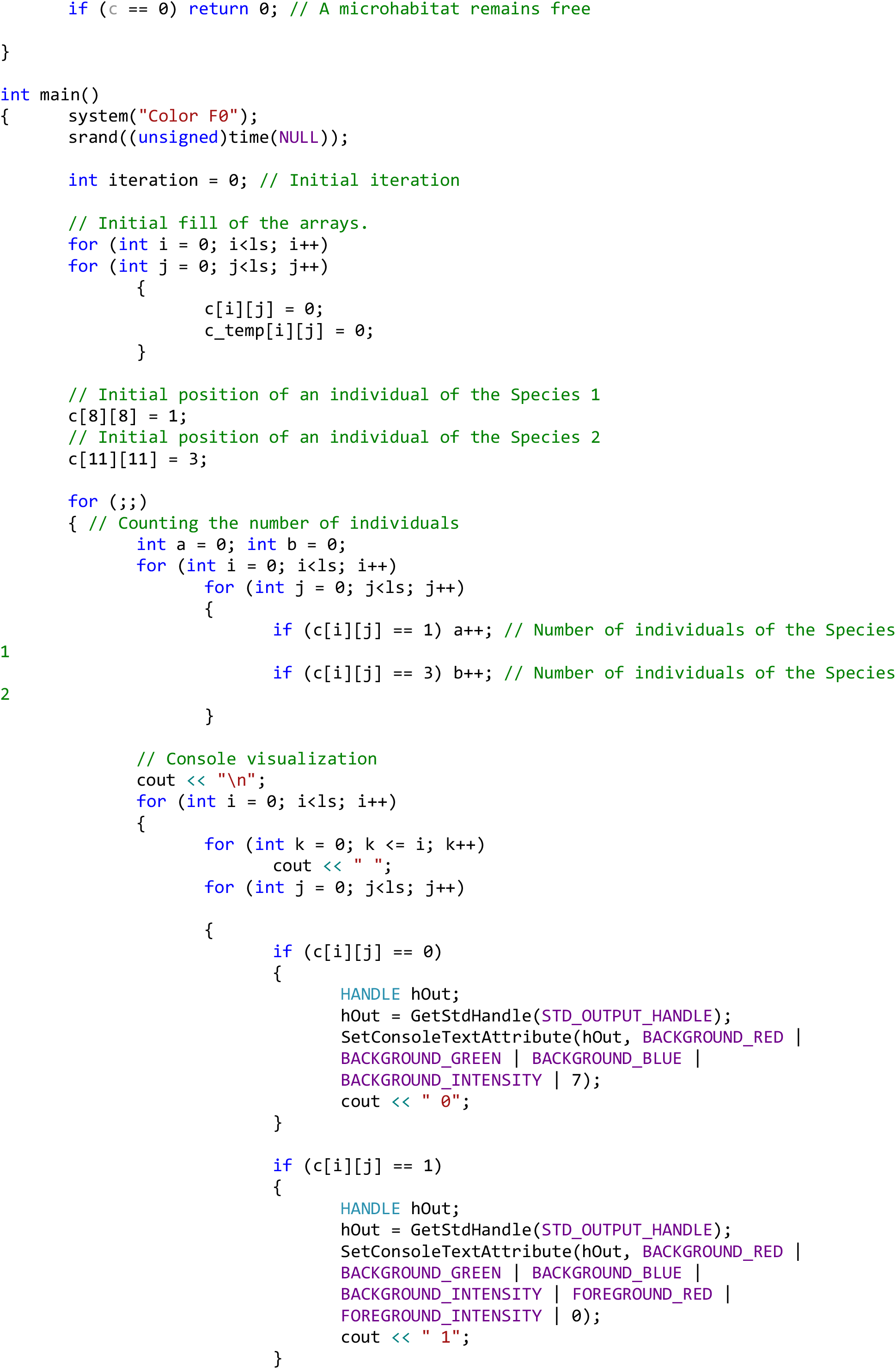
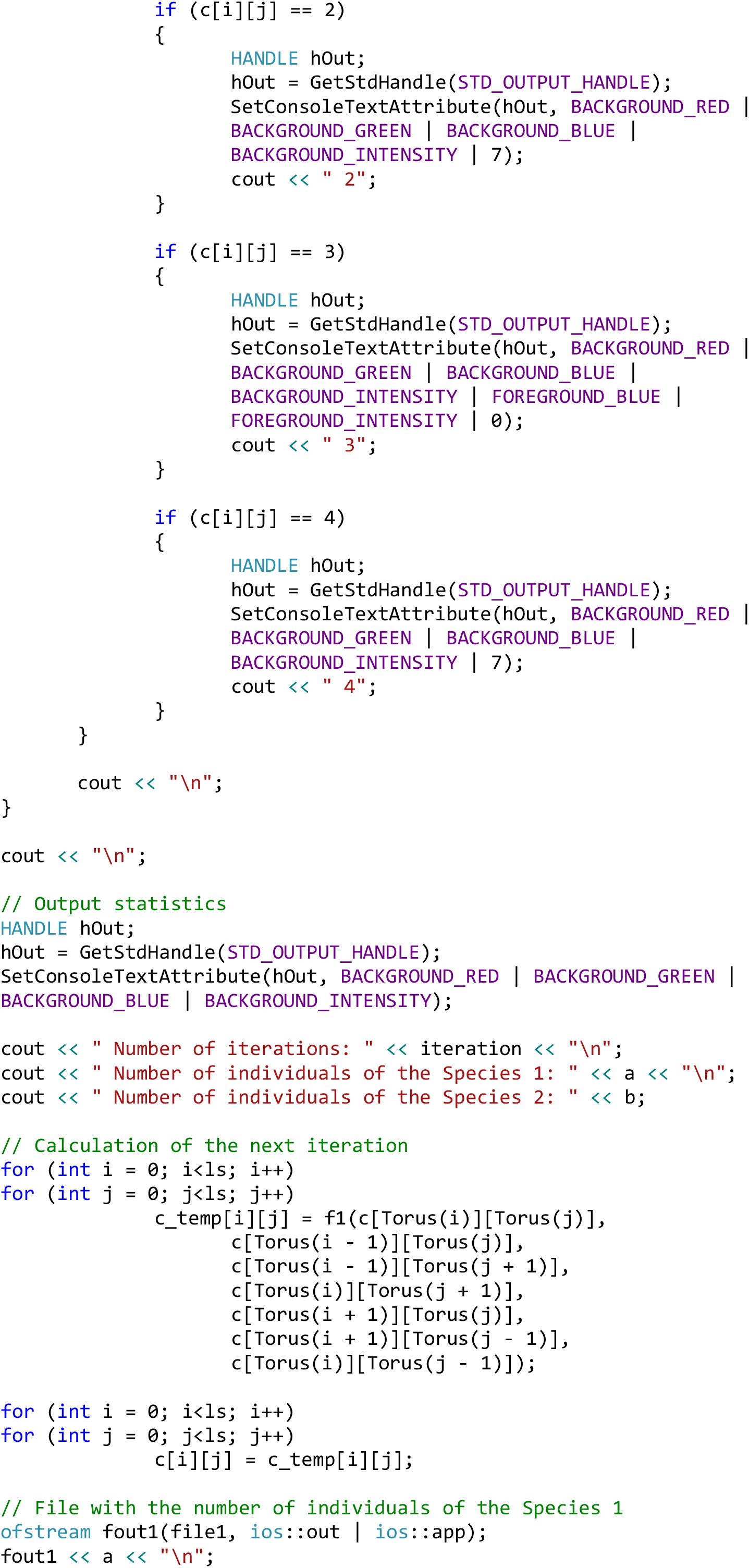
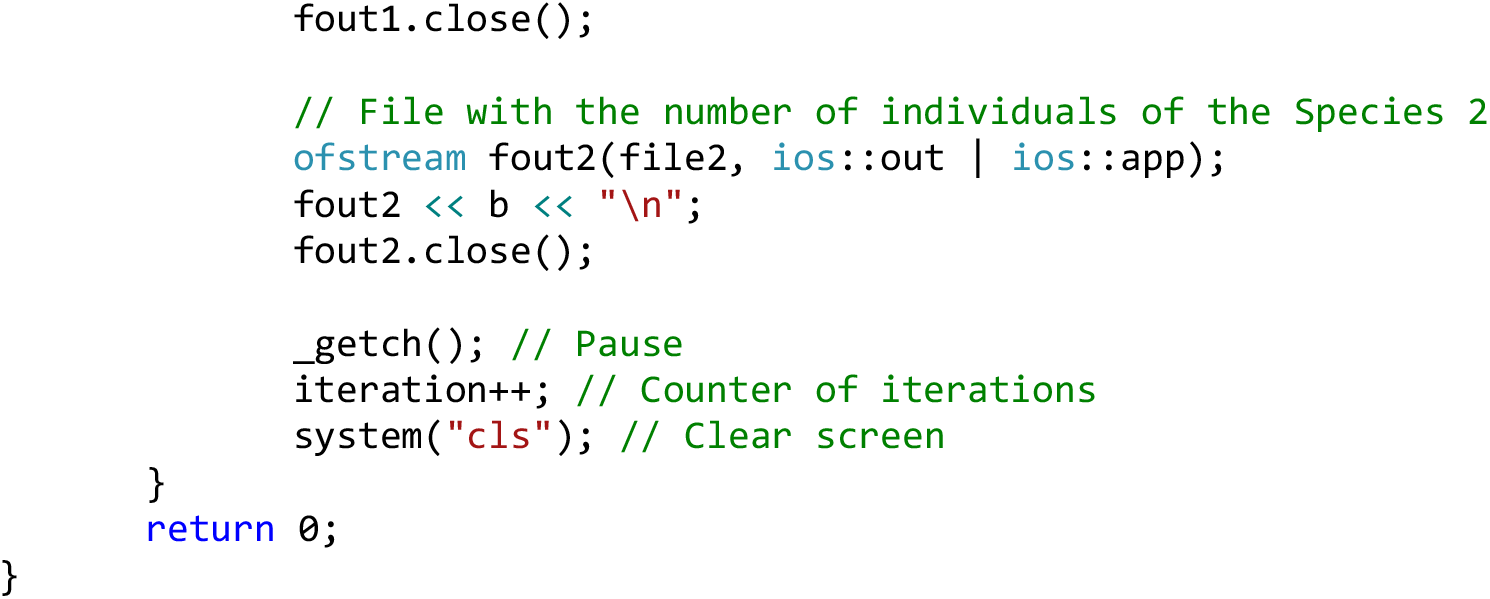
Model 2. Coexistence of complete competitors with the initial 10% difference in fitness and with dependence of the fitness on a number of locally competing individuals. The initial pattern of the habitat is populated by two single individuals of two competing species. The increase in the number of individuals of a species, locally competing for reproduction in a given free microhabitat by one unit, increases its fitness by 0.08 (by 8%).

**Source code 4.**
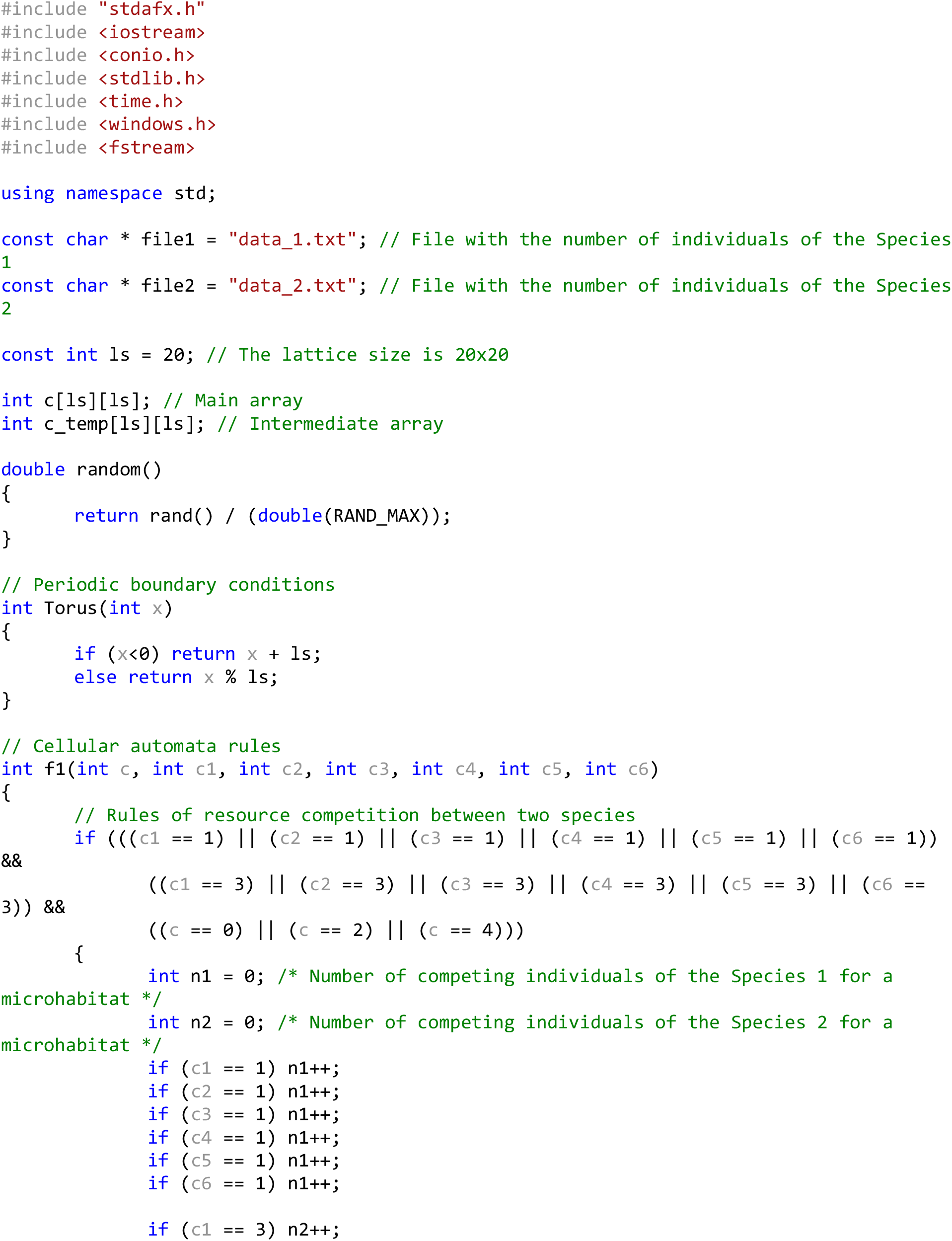
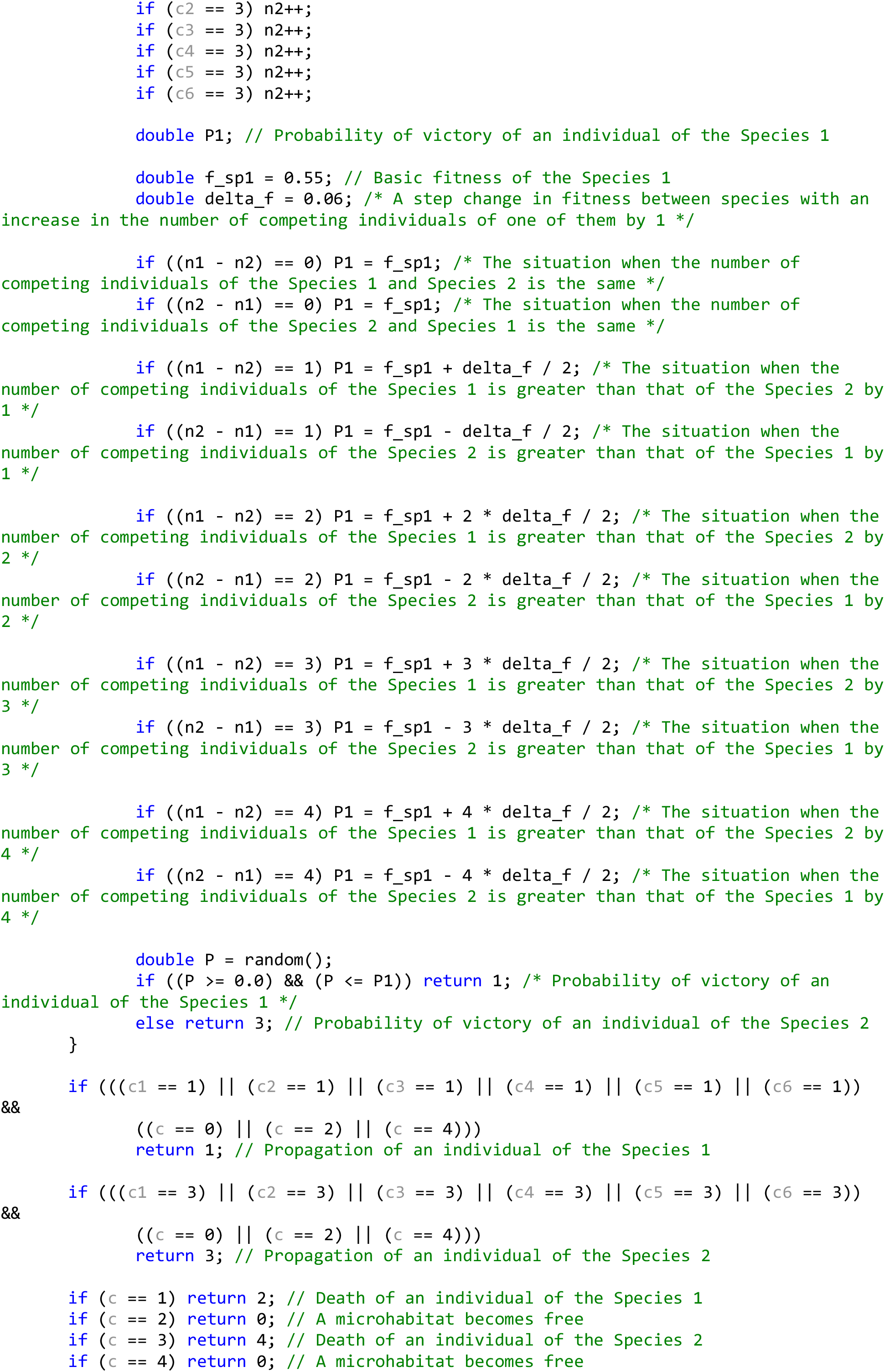
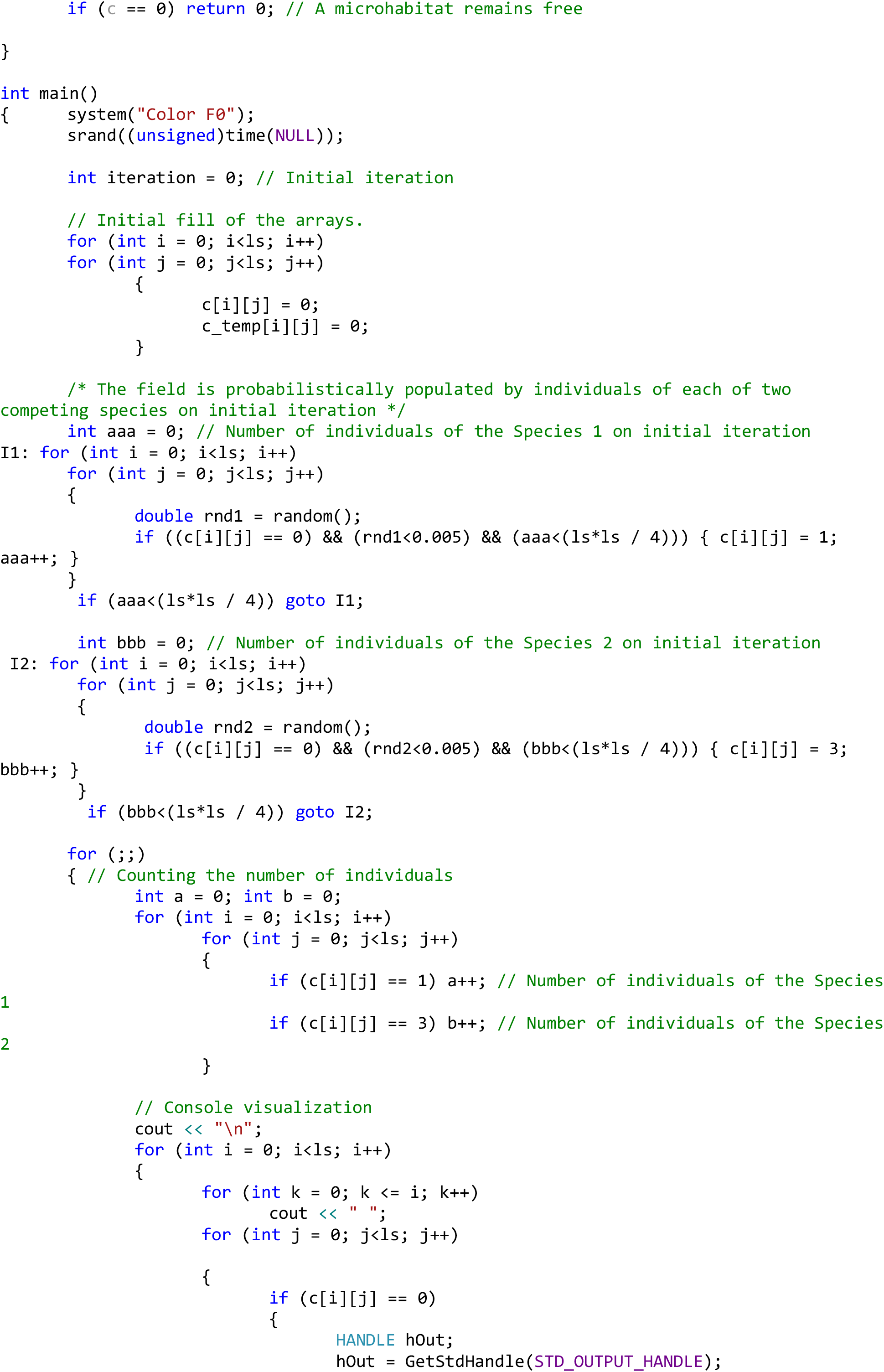
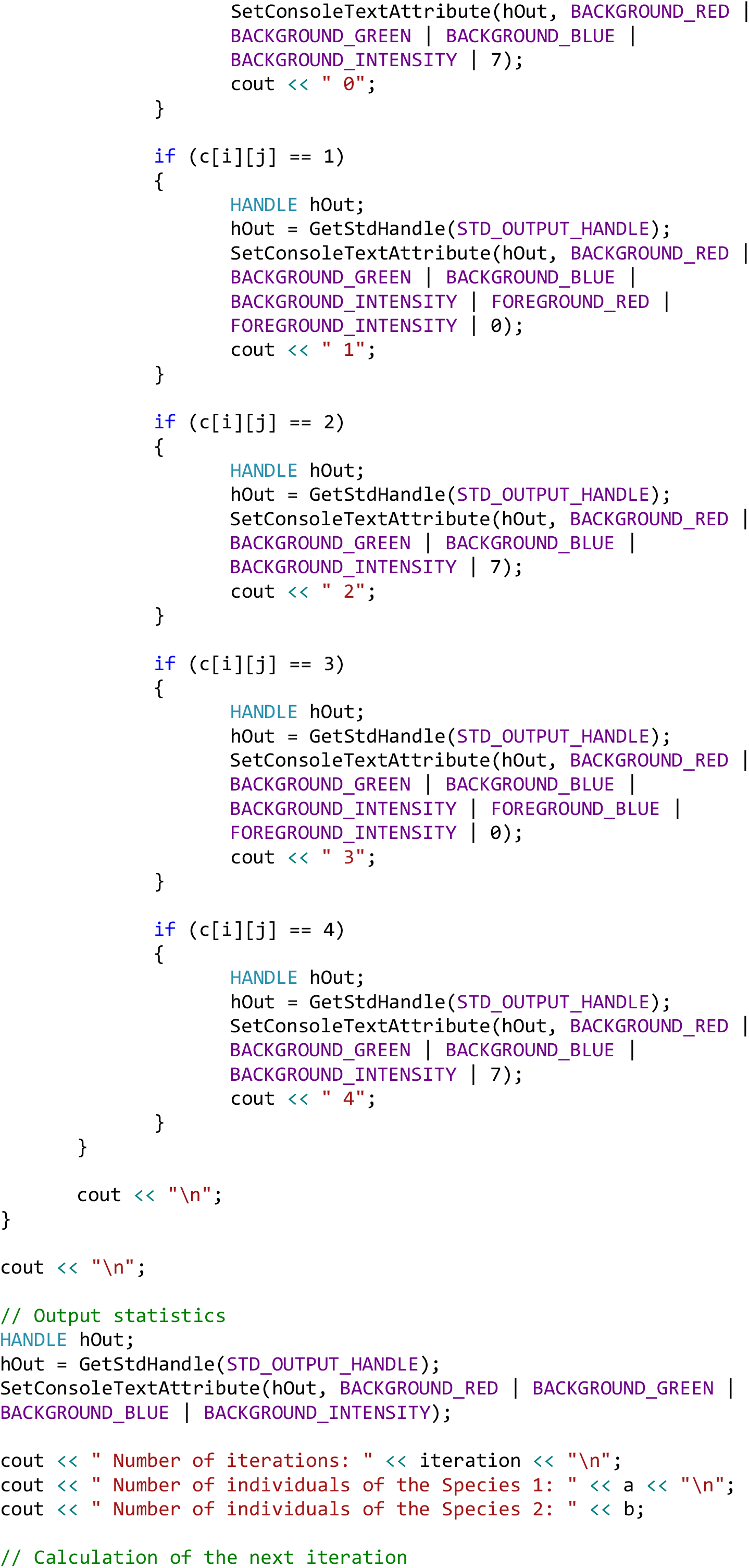
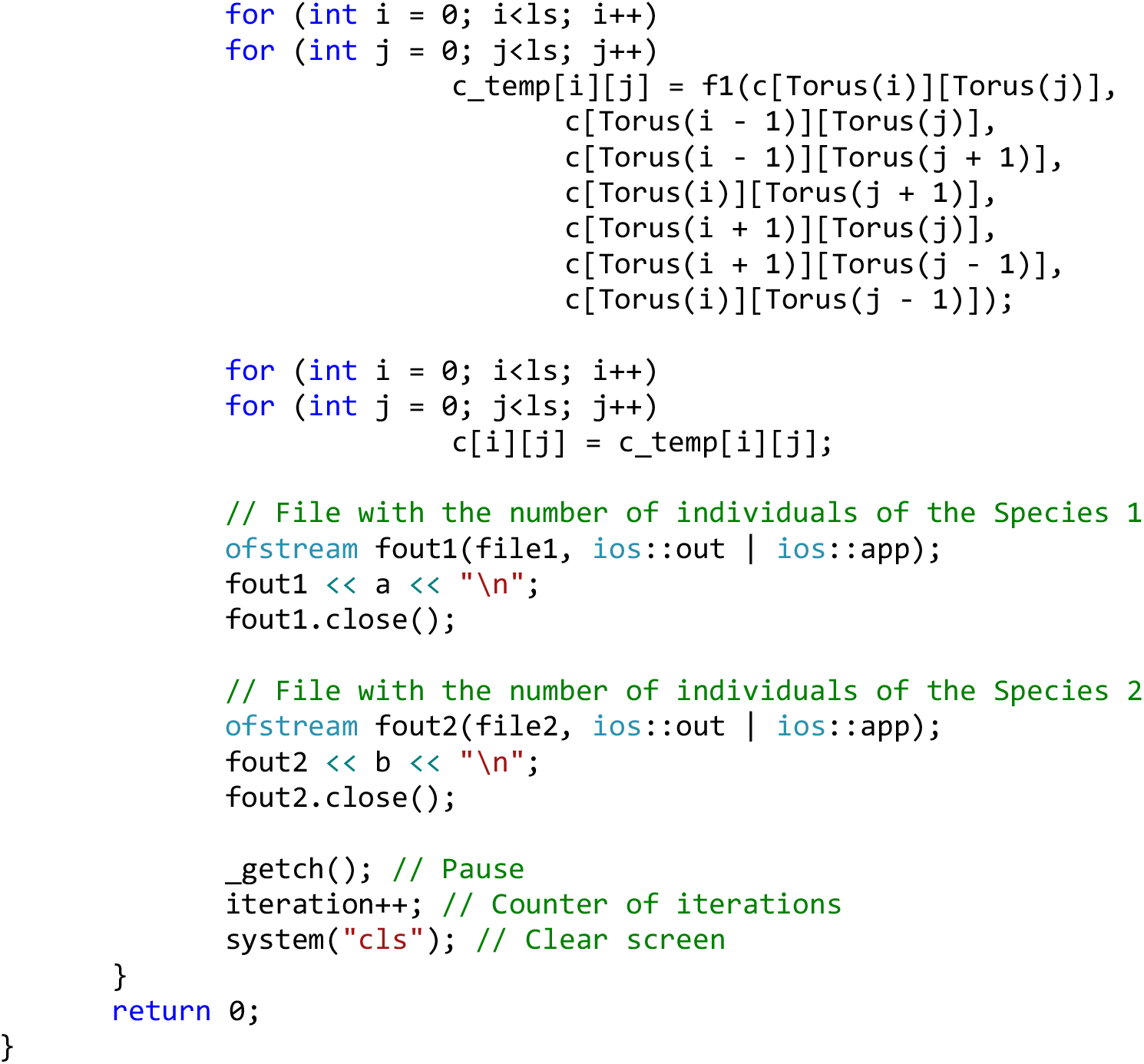
Model 2. Coexistence of complete competitors with the initial 10% difference in fitness and with dependence of the fitness on a number of locally competing individuals. The initial pattern of the habitat is randomly populated by individuals - 25% by the Species 1, 25% by the Species 2 and 50% of the habitat remains free. The increase in the number of individuals of a species, locally competing for reproduction in a given free microhabitat, by one unit, increases its fitness by 0.06 (by 6%).

## Notes

https://figshare.com/projects/Raw_Data_of_the_manuscript_Axiomatic-deductive_theory_of_competition_of_complete_competitors_coexistence_exclusion_and_neutrality_in_one_model_/20840

### Competing Interest Statement

The authors have declared no competing interest.

